# The tumor microenvironment drives transcriptional phenotypes and their plasticity in metastatic pancreatic cancer

**DOI:** 10.1101/2020.08.25.256214

**Authors:** Srivatsan Raghavan, Peter S. Winter, Andrew W. Navia, Hannah L. Williams, Alan DenAdel, Radha L. Kalekar, Jennyfer Galvez-Reyes, Kristen E. Lowder, Nolawit Mulugeta, Manisha S. Raghavan, Ashir A. Borah, Kevin S. Kapner, Sara A. Väyrynen, Andressa Dias Costa, Raymond W.S. Ng, Junning Wang, Emma Reilly, Dorisanne Y. Ragon, Lauren K. Brais, Alex M. Jaeger, Liam F. Spurr, Yvonne Y. Li, Andrew D. Cherniack, Isaac Wakiro, Asaf Rotem, Bruce E. Johnson, James M. McFarland, Ewa T. Sicinska, Tyler E. Jacks, Thomas E. Clancy, Kimberly Perez, Douglas A. Rubinson, Kimmie Ng, James M. Cleary, Lorin Crawford, Scott R. Manalis, Jonathan A. Nowak, Brian M. Wolpin, William C. Hahn, Andrew J. Aguirre, Alex K. Shalek

**Affiliations:** Department of Medical Oncology, Dana-Farber Cancer Institute, Boston, MA; Broad Institute of MIT and Harvard, Cambridge, MA; Harvard Medical School, Boston, MA; Department of Medicine, Brigham and Women’s Hospital, Boston, MA; Institute for Medical Engineering & Science, Massachusetts Institute of Technology, Cambridge, MA; Koch Institute for Integrative Cancer Research, Massachusetts Institute of Technology, Cambridge, MA; Department of Chemistry, Massachusetts Institute of Technology, Cambridge, MA; Center for Computational Molecular Biology, Brown University, Providence, RI; Division of Applied Mathematics, Brown University, Providence, RI; Center for Cancer Genomics, Dana-Farber Cancer Institute, Boston, MA; Department of Oncologic Pathology, Dana-Farber Cancer Institute, Boston, MA; Department of Surgery, Brigham and Women’s Hospital, Boston, MA; Department of Biostatistics, Brown University, Providence, RI; Microsoft Research New England, Cambridge, MA; Department of Biological Engineering, Massachusetts Institute of Technology, Cambridge, MA; Department of Pathology, Brigham and Women’s Hospital, Boston, MA; Ragon Institute of MGH, MIT, and Harvard, Cambridge, MA

**Keywords:** Pancreatic cancer, Single-cell RNA sequencing, Cancer cell states, Transcriptional phenotypes, Plasticity, Patient-derived organoid models, Genotype-phenotype relationships, Tumor microenvironment, Liver metastases, Tumor heterogeneity

## Abstract

Bulk transcriptomic studies have defined classical and basal-like gene expression subtypes in pancreatic ductal adenocarcinoma (PDAC) that correlate with survival and response to chemotherapy; however, the underlying mechanisms that govern these subtypes and their heterogeneity remain elusive. Here, we performed single-cell RNA-sequencing of 23 metastatic PDAC needle biopsies and matched organoid models to understand how tumor cell-intrinsic features and extrinsic factors in the tumor microenvironment (TME) shape PDAC cancer cell phenotypes. We identify a novel cancer cell state that co-expresses basal-like and classical signatures, demonstrates upregulation of developmental and KRAS-driven gene expression programs, and represents a transitional intermediate between the basal-like and classical poles. Further, we observe structure to the metastatic TME supporting a model whereby reciprocal intercellular signaling shapes the local microenvironment and influences cancer cell transcriptional subtypes. In organoid culture, we find that transcriptional phenotypes are plastic and strongly skew toward the classical expression state, irrespective of genotype. Moreover, we show that patient-relevant transcriptional heterogeneity can be rescued by supplementing organoid media with factors found in the TME in a subtype-specific manner. Collectively, our study demonstrates that distinct microenvironmental signals are critical regulators of clinically relevant PDAC transcriptional states and their plasticity, identifies the necessity for considering the TME in cancer modeling efforts, and provides a generalizable approach for delineating the cell-intrinsic versus -extrinsic factors that govern tumor cell phenotypes.

## INTRODUCTION

Classification of human malignancies by genotype has provided important insights into tumor biology as well as a framework to guide therapeutic selection in many cancers (Hyman et al., 2017). However, tumors also exhibit clinically relevant transcriptional variation that can influence malignant progression and therapeutic response. The application of single-cell RNA-sequencing (scRNA-seq) to tumor specimens has afforded a means to characterize the malignant and non-malignant cellular components of the tumor microenvironment (TME) and their heterogeneity at unprecedented resolution (Kim et al., 2018; Patel et al., 2014; Puram et al., 2017; Sade-Feldman et al., 2019; Suva and Tirosh, 2019; Tirosh et al., 2016a; van Galen et al., 2019; Venteicher et al., 2017). These analytical approaches have further enabled the re-examination of existing transcriptional taxonomies, revealing structured heterogeneity within malignant populations and reframing our understanding of bulk measurements in multiple cancers (Filbin et al., 2018; Hovestadt et al., 2019; Neftel et al., 2019; Patel et al., 2014; Tirosh et al., 2016b; Venteicher et al., 2017).

The phenotypic variability observed in human tumors often reflects the underlying cancer cell genetics. Specific mutations can program cancer cell states and, in some cases, serve as biomarkers for treatment (Filbin et al., 2018; Hovestadt et al., 2019; van Galen et al., 2019; Venteicher et al., 2017). Yet, in other instances, transcriptional phenotypes are not strongly associated with specific mutational patterns (Nam et al., 2021). In these tumors, cell-extrinsic TME interactions may influence malignant cellular attributes, but our understanding of reciprocal signaling between malignant cells and the TME is rudimentary. Mapping the cell-intrinsic and -extrinsic factors that impact tumor cell states and determining which ones drive phenotypic transitions would yield important insights into the biologic basis for clinical disease phenotypes and drug resistance.

For pancreatic ductal adenocarcinoma (PDAC), bulk RNA-seq profiling has defined two major transcriptional programs, basal-like/squamous (hereafter referred to as basal) and classical. The basal subtype is strongly associated with a poorer prognosis and greater treatment resistance (Aguirre et al., 2018; Aung et al., 2018; Bailey et al., 2016; Cancer Genome Atlas Research Network, 2017; Chan-Seng-Yue et al., 2020; Collisson et al., 2011; Connor et al., 2019; Moffitt et al., 2015; O’Kane et al., 2020; Porter et al., 2019; Tiriac et al., 2018), but the roles of cell-intrinsic and -extrinsic factors in determining these cell states and their sensitivity to different therapies are not well understood. A limited number of genomic alterations, including *TP53* mutational status and *c-MYC* or *KRAS* amplifications, have been associated with the more therapy-resistant basal state (Bailey et al., 2016; Cancer Genome Atlas Research Network, 2017; Chan-Seng-Yue et al., 2020; Hayashi et al., 2020; Schleger et al., 2002). Recent studies have also suggested that high levels of KRAS expression and signaling can induce the basal state, but others have demonstrated that basal PDAC cells exhibit RAS-independence (Chan-Seng-Yue et al., 2020; Collisson et al., 2011; Miyabayashi et al., 2020; Muzumdar et al., 2017). These findings suggest that while genomic activation of *KRAS* plays an important role in oncogenesis, other non-genetic and microenvironmental factors may also be critical in regulating downstream cellular states.

Although the majority of patients with PDAC present with and succumb to metastatic disease (Siegel et al., 2020), our current understanding of PDAC is largely derived from resected primary tumors (Aguirre et al., 2018; Cancer Genome Atlas Research Network, 2017; Siegel et al., 2020). While several recent studies have described the desmoplastic stromal microenvironment and immune infiltration in primary PDAC (Balachandran et al., 2019; Bernard et al., 2019; Biffi et al., 2019; Elyada et al., 2019; Ligorio et al., 2019; Ohlund et al., 2017), we lack a detailed characterization of the immune and stromal cells that constitute metastatic PDAC lesions. The local TME in the pancreas is likely different from metastatic sites in other organs (Ho et al., 2020), and given the strong association of transcriptional subtype with survival and drug resistance (Aguirre et al., 2018; Aung et al., 2018; Bailey et al., 2016; Cancer Genome Atlas Research Network, 2017; Chan-Seng-Yue et al., 2020; Collisson et al., 2011; Connor et al., 2019; Moffitt et al., 2015; O’Kane et al., 2020; Porter et al., 2019; Tiriac et al., 2018), understanding whether specific inputs from the metastatic niche can specify transcriptional phenotype is of great importance to targeting therapeutic resistance in PDAC.

To better understand the interplay between genetics, transcriptional state, and the metastatic TME, we developed and employed an optimized translational workflow to perform both high-resolution profiling of PDAC patient tissue using scRNA-seq (Gierahn et al., 2017; Hughes et al., 2020) and derivation of matched organoid models (Boj et al., 2015; Tiriac et al., 2018) from the same metastatic core needle biopsy. Using matched *in vivo* observations and *ex vivo* experimental studies, we describe a tumor cell atlas of metastatic PDAC, identify a new intermediate transitional PDAC cancer cell state, uncover distinct site- and subtype-specific TMEs, and demonstrate that microenvironmental signals are critical regulators of transcriptional subtypes and their plasticity.

## RESULTS

### A clinical pipeline for matched single-cell profiling and organoid model generation

We established a pipeline for collecting core needle biopsies from patients with metastatic PDAC (n=23) to generate matched scRNA-seq profiles and organoid models (**Figure 1A**; **Supplemental Figure S1A**; **Supplemental Table S1**). Most samples were obtained from metastatic lesions residing in the liver (19/23), and the majority (21/23) were analyzed by targeted DNA-sequencing, yielding the expected mutational pattern for this disease (**Figure 1B**) (Aguirre et al., 2018; Bailey et al., 2016; Cancer Genome Atlas Research Network, 2017). Our pipeline generated approximately 1,000 high-quality single cells per biopsy (n=23,042 total cells) and successful early-passage organoid cultures from 70% of patient tumor samples (16/23 samples reaching at least passage 2; **Figure 1B**; **Supplemental Figure S1A,B**). Dimensionality reduction and shared nearest neighbor (SNN) clustering of the biopsy cells revealed substantial heterogeneity at the single-cell level (**Supplemental Figure S1C,D**; **Methods**). Consistent with other studies of human cancer, we observed patient-specific and admixed clusters of single cells suggesting the presence of both malignant and non-malignant cells in each biopsy (Kim et al., 2018; Puram et al., 2017; Sade-Feldman et al., 2019; Tirosh et al., 2016a; van Galen et al., 2019). To confirm which clusters were comprised of malignant cells, we inferred transcriptome-wide CNVs from our single-cell data as previously described (Patel et al., 2014; Tirosh et al., 2016b). CNV alteration scores separated putative cancerous and non-cancerous cells in each biopsy and demonstrated high concordance with reference targeted DNA-seq (**Figure 1C**; **Supplemental Figure S1E,F**). CNV analysis paired with manual inspection of expression patterns for known markers across single cells supported the identification of cancerous cells as well as 11 unique non-cancerous cell types (**Figure 1D,E**; **Supplemental Figure S1D-I**; **Supplemental Table S2**). Thus, we established a robust workflow capable of recovering high quality malignant (n=7,740) and non-malignant (n=15,302) populations from metastatic PDAC needle biopsies while also enabling simultaneous generation of matched organoid models.

**Figure 1.**
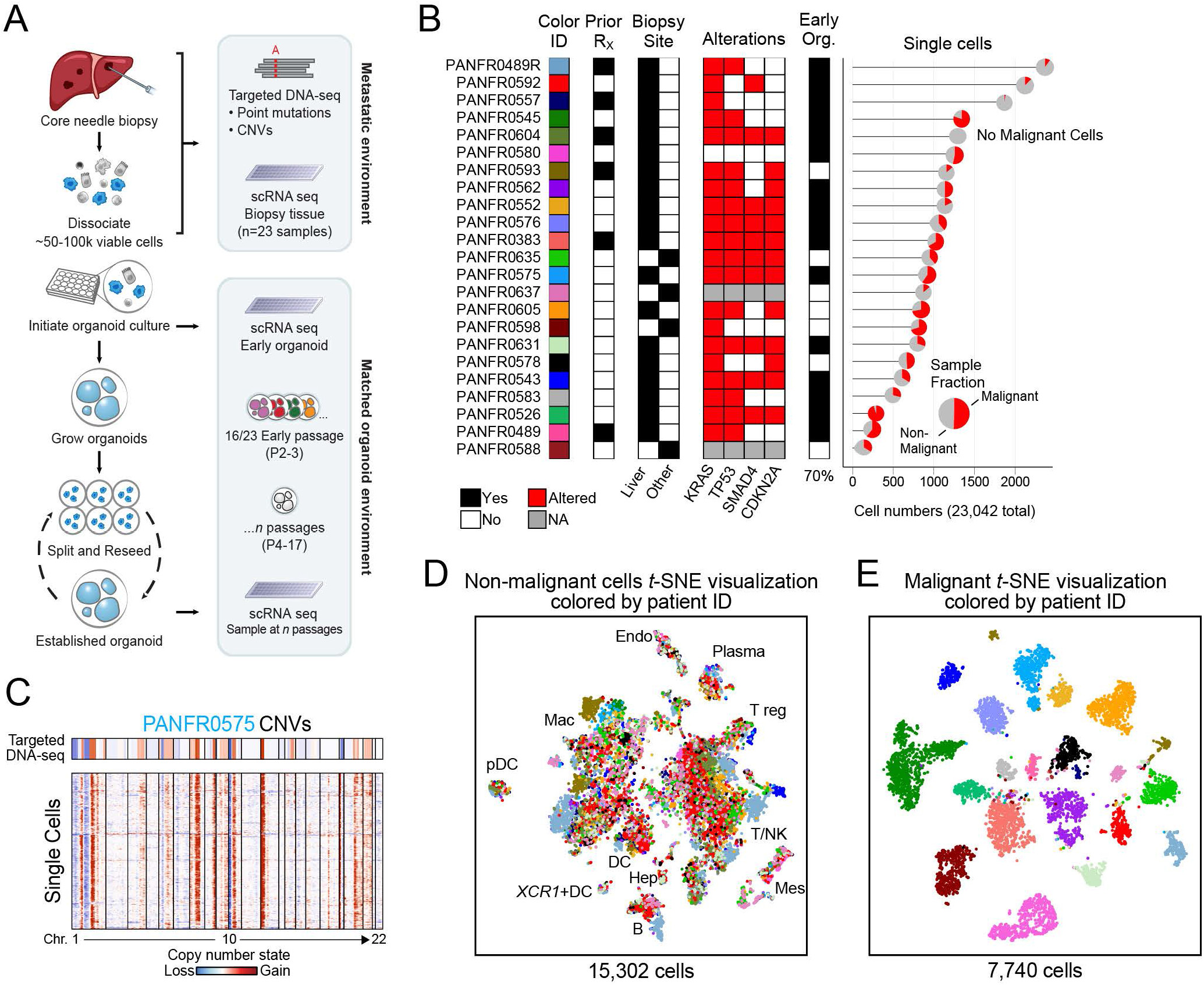
A clinical pipeline for matched single-cell RNA-seq and organoid generation from metastatic PDAC biopsies. **(A)** Pipeline for collecting patient samples, and dissociation and allocation for scRNA-seq and parallel organoid development. **(B)** Clinical and molecular features for all patients included in the dataset (Rx = Therapy; Other = Adrenal (PANFR0637), Omentum (PANFR0635, PANFR0598), Peritoneum (PANFR0588); Org. at P2 = Organoid measured at passage 2). Mutations were determined by bulk targeted DNA-seq (Red, Altered; White, wildtype; Grey, Data not available). Number of single cells captured per biopsy and their malignant and non-malignant fraction is visualized at the right. **(C)** Example bulk targeted DNA-seq (top) and single-cell inferred CNV profiles (rows, bottom) arranged by chromosome (columns) from PANFR0575. **(D-E)** *t*-distributed stochastic neighbor embedding (*t*-SNE) visualization for non-malignant (**D**) and malignant (**E**) single cells in the biopsy cohort. Cells are colored by patient as in **B**. Endo, Endothelial; Mes, Mesenchymal; B, B-cell; Hep, Hepatocyte; DC, Dendritic cell; pDC, Plasmacytoid dendritic cell; Mac, Macrophage; T, T-cell; NK, Natural killer cell. *See also Supplemental Figures S1 & S2; Supplemental Tables S1 & S2*.

### Tumor cell transcriptional subtypes in metastatic PDAC include an intermediate transitional state

We applied principal component analysis (PCA) to examine transcriptional variation across cancer cells from all biopsy samples. CNV-altered cells from one biopsy, PANFR0580, separated from the rest of the samples (**Figure 1B**; **Supplemental Figure S2A**). Based on expression of known neuroendocrine markers (*TTR, CHGA*) and subsequent pathology review we reclassified this sample as a pancreatic neuroendocrine tumor (PanNET) and used it as a non-PDAC reference cell population. Among the remaining 7,078 PDAC cells, we found that genes driving the first 3 PCs were enriched for signatures of epithelial/mesenchymal transition [EMT, PC1, (Groger et al., 2012)], basal/classical state [PC2, (Moffitt et al., 2015)], and cell cycle [PC3, (Tirosh et al., 2016a)] (**Supplemental Figure S2B**). When we scored all malignant cells within our cohort for basal and classical gene expression, we observed that they inhabited a graded continuum of expression states from strongly basal to strongly classical (**Figure 2A**). Correlation analysis across malignant cells revealed 1,909 genes significantly associated with either basal or classical expression scores (**Supplemental Figure S2C**; **Supplemental Table S3**; **Methods**). Inspection of these genes revealed that basal cells are defined by squamous and mesenchymal features and co-express programs associated with transforming growth factor beta (TGF-β) signaling, WNT signaling, and cell cycle progression (Groger et al., 2012; Kim et al., 2017; Tirosh et al., 2016a). Conversely, epithelial and pancreatic lineage programs are enriched in classical subtype PDAC cells (**Supplemental Figure S2D,E**).

**Figure 2.**
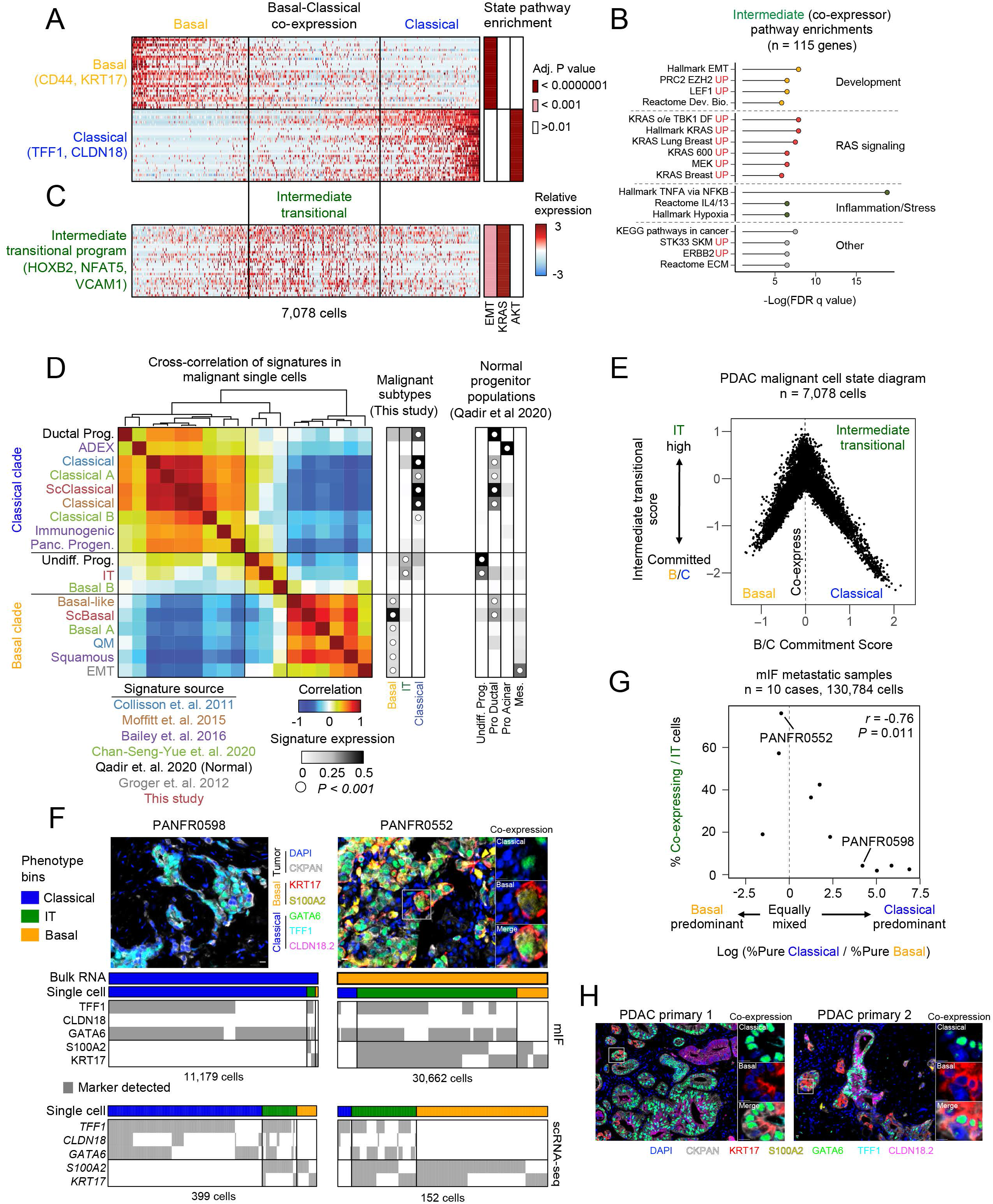
An intermediate transitional state bridges basal and classical phenotypes. **(A)** Heatmaps depict the expression of basal and classical expression programs and highlight the co-expressing intermediates (n=30 genes each). **(B)** Gene set enrichment analysis for the 115 genes specific to the co-expressing intermediate state. **(C)** The intermediate transitional (“IT”) expression program (n = 30 genes) is enriched by co-expressing cells. Enrichment adjusted *P*-values (hypergeometric test) for EMT, *KRAS*, and *AKT* gene sets are indicated at right for each gene expression program in **A** and **C**. **(D)** Cross-correlation between new and previously proposed expression signatures (rows and columns; text color = source, below) in our PDAC single-cells. Average expression for each signature (rows) is shown at the right for cells in the malignant subtypes from our cohort and the normal pancreatic progenitor cells from Qadir et al., 2020. White dot indicates the subset with the highest average significant expression for each signature (Kruskal-Wallis test); no white dot indicates no significant expression. **(E)** Malignant cell state diagram for PDAC. Basal-classical commitment score (x axis) and IT score (y axis) for all 7,078 malignant cells (**Methods**). **(F)** Multiplex immunofluorescence analysis (mIF) identifies co-expressing IT cells in matched metastatic samples. Top are representative images from two cases (white box indicates region for co-expression insets at right), and bottom indicates marker detection patterns for mIF and matched scRNA-seq data (**Methods**). Scale bar represents 10 μm. **(G)** Frequency of co-expressing IT cells is correlated with balanced representation of pure basal and classical phenotypes by mIF within individual samples. Log ratio of % basal and classical cells in each sample (x axis) versus their % co-expressing / IT cells (y axis). **(H)** Co-expressing IT cells are also identified in primary PDAC samples by mIF. Scale bar represents 10μm. *See also Supplemental Figures S2, S3 & S4; Supplemental Tables S3 & S4.*

Strikingly, we observed that the basal and classical programs were not mutually exclusive; rather, we identified a large population of cells that co-expressed features of both programs to varying degrees (**Figure 2A**; **Supplemental Figure S3A,B**). In developmental contexts, cell state commitment is often a continuous process where mixing/co-expression of state markers indicates state transitions (Nam et al., 2021). Similarly, the large fraction of intermediate co-expressing cells identified in our single-cell snapshots suggests state transitions may be an ongoing and frequent process in human PDAC tumors. We identified 115 genes whose expression was correlated with this co-expressor intermediate state and enriched for developmental, Ras signaling, and inflammation/stress response gene sets (**Figure 2B**; **Supplemental Figure S3C,D**; **Supplemental Table S3**; **Methods**). Signatures of RAS signaling were enriched in the intermediate state even compared with basal and classical programs, and, by contrast, classical phenotypes were enriched for Akt-associated gene sets and showed little evidence of EMT or RAS enrichments (**Figure 2A-B**; **Supplemental Figure S2E**).

Since this intermediate signature showed enrichment for developmental gene programs, we next assessed whether this signature overlapped with any phenotypes recently reported in the normal pancreas progenitor niche (Qadir et al., 2020). We found that both basal and classical gene expression signatures were expressed by pro-ductal progenitor cells, while the intermediate gene expression program was enriched in an undifferentiated, stress-responsive progenitor population (**Supplemental Figure S3E,F**) (Qadir et al., 2020). Thus, based on its enrichment for developmental and stress-responsive gene sets, overlap with populations in the normal progenitor niche, and co-expression of basal and classical programs suggestive of a transitional state, we termed this phenotype “Intermediate transitional” (IT) (**Figure 2C**).

To further contextualize this cell state, we compared signatures proposed by prior bulk RNA-sequencing studies to clarify potential inter-relationships (Aguirre et al., 2018; Bailey et al., 2016; Chan-Seng-Yue et al., 2020; Collisson et al., 2011; Moffitt et al., 2015). Pairwise correlation of all established signatures in malignant cells revealed that many contribute overlapping information and reflect similar underlying biology within either basal or classical clades, but that the IT signature is unique and not well described by established bulk RNA-seq signatures (**Figure 2D**). Taken together, these findings suggest that malignant PDAC cells organize in a tripartite cell state framework that spans committed basal and classical phenotypes, with considerable signature co-expression in single cells (**Figure 2E**). Similar to the variation in EMT scores observed in basal tumor cells (**Supplemental Figure S3A**) (Chan-Seng-Yue et al., 2020; Connor et al., 2019), we noted heterogeneity among co-expressing cells for the IT program.

### Multiplex immunofluorescence confirms co-expressing IT cells in metastatic and primary PDAC

To compare to bulk RNA-seq studies, we clustered pseudo-bulk averages of the malignant cells from each biopsy and observed separation of tumors into those that expressed predominantly basal, classical, or IT signatures (**Supplemental Figure S3G-I**). However, individual tumors exhibited significant heterogeneity at the cellular level, with mixing of malignant cell populations expressing at least two and frequently all three cell states within the same patient specimen (**Supplemental Figure S3J**). To validate the extensive heterogeneity and the presence of co-expressing IT cells in our metastatic cohort, we used a subtype-specific multiplex immunofluorescence (mIF) panel to categorize single tumor cells by their patterns of marker detection in 10 matched cases from our single cell study (**Supplemental Figure S4A**; **Supplemental Table S4**; **Methods**). We observed extensive overlap of basal and classical markers within single cells at the protein level, corroborating the existence of co-expressing IT cells using an orthogonal method (**Figure 2F**; **Supplemental Figure S4B**). Encouragingly, we observed significant correlation within subtype (average *r =* 0.52) as compared to between subtypes (average *r =* 0.06, *P* < 10^−7^, Student’s T test) using this orthogonal method. We also observed high concordance between the two methods, giving confidence that we are accurately sampling the distribution of states present in each sample (average *r* = 0.45; **Supplemental Figure S4C**, white dots). As with scRNA-seq, we observed mixing of basal, classical, and IT cells within individual patient specimens by mIF subtyping. The frequency of co-expressing cells was correlated with balanced representation of pure basal and classical phenotypes within individual samples, consistent with the co-expressing IT phenotype as a transitional state (**Figure 2G**). Indeed, none of the tumors evaluated with mIF contained a mix of pure basal and classical phenotypes in the absence of co-expressing IT cells. We also identified co-expressing cells in primary tumor samples which suggests that IT phenotypes may be a general feature of PDAC tumors in both the localized and metastatic settings (**Figure 2H**; **Supplemental Figure S4D**).

### Microenvironment is dominant to *KRAS* amplifications in determining transcriptional subtype

We next searched for potential molecular regulators of the observed tumor cell transcriptional heterogeneity. In PDAC, the vast majority of tumors harbor clonal *KRAS* point mutations, including all of the PDAC samples in our cohort (**Figure 1B**). While point mutations in *KRAS* do not appear to determine transcriptional subtype, several studies have suggested that amplifications in *KRAS* associate with more basal features (Chan-Seng-Yue et al., 2020; Miyabayashi et al., 2020), while amplifications of lineage transcription factors like *GATA6* associate with classical phenotypes (Chan-Seng-Yue et al., 2020). To assess for such genotype-phenotype relationships in our single-cell cohort, we inferred copy number variation for common PDAC alterations (*KRAS*, *TP53*, *SMAD4*, and *CDKN2A*) and lineage-associated transcription factors (*HNF4G* and *GATA6*) from scRNA-seq expression data using a previously established Hidden Markov model workflow (**Methods**) (Fan et al., 2018; Patel et al., 2014; Tirosh et al., 2016b). Encouragingly, we observed a significant association between single-cell inferred *KRAS* copy number gain and basal phenotypes (*P<0.03* Fisher’s exact test), and also between inferred *CDKN2A* copy loss and IT phenotypes (*P<0.003* Fisher’s exact test; **Figure 3A**). While we found that cells derived from samples with inferred *KRAS* amplifications had a strong preference for the basal subtype, these cells could still span all three phenotypic categories or be predominantly classical within an individual tumor (**Figure 3B-D)**.

**Figure 3.**
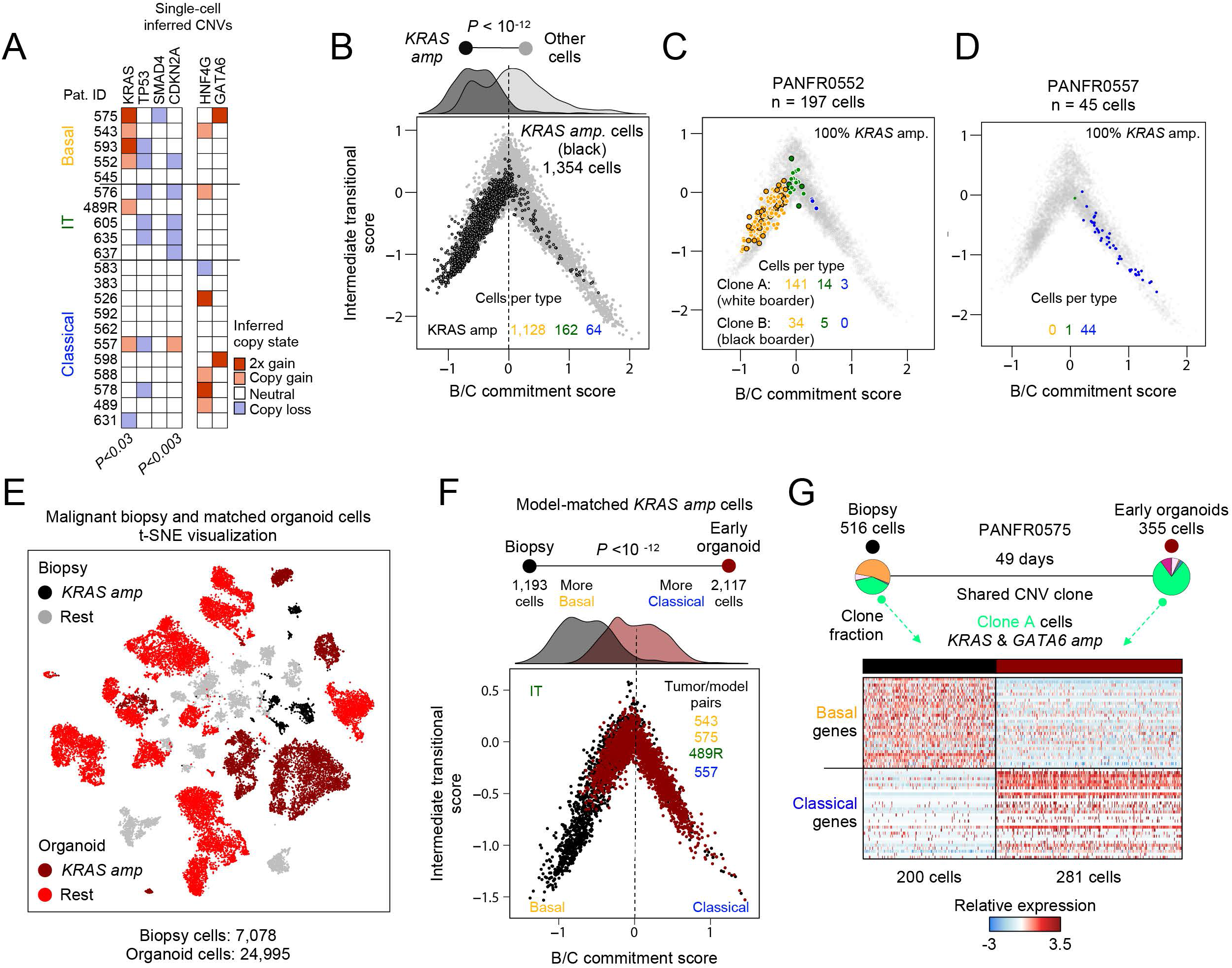
Microenvironment dictates phenotype in *KRAS*-amplified tumor cells. **(A)** Single-cell inferred copy number alterations for each sample in the biopsy cohort (**Methods**). Tumors are grouped by expression of their dominant subtype based on the clustering in **Supp. Fig. S3G**, *P*-values comparing presence of each alteration among the groups (Basal, Classical, IT) are determined by Fisher’s exact test. **(B)** Malignant cell state diagram as in Figure 2E but highlighting all *in vivo KRAS*-amplified tumor cells (black border) across the states. **(C)** Similar to **B**, but highlighting PANFR0552 *KRAS*-amplified malignant cell heterogeneity. White and black borders correspond to separate CNV sub-clones (both *KRAS* amplified) and color fill denotes transcriptional subtype. **(D)** Similar to **B**, but highlighting PANFR0557 *KRAS*-amplified malignant cell heterogeneity. Color fill denotes transcriptional subtype. **(E)** *t*-SNE visualization of all biopsy (grey and black) and matched organoid cells (red and dark red) from iterative passages. *KRAS*-amplified tumor cells from *in vivo* specimens (black) and organoid models (dark red) are highlighted with distinct colors. **(F)** Cell state diagram for all cells with inferred *KRAS* amplifications in biopsy (grey) and organoid (red) microenvironments. *P*-value compares biopsy versus early passage organoid score distributions (top density) and was determined by student’s T test. **(G)** Clonal fractions (pie charts) from the *KRAS*-amplified PANFR0575 sample in biopsy and organoid conditions. Heatmap shows the relative expression in single cells from plastic clone A (bright green) in both conditions.

To further examine this genotype-to-phenotype association, we tested if *KRAS* amplification was sufficient to specify the basal phenotype in an *ex vivo* environment. We initiated patient-derived organoid cultures from matched PDAC biopsies and serially sampled them over time with scRNA-seq (**Figure 1A**). CNV-confirmed “early” organoid cells (first passage measured, n=2,117 cells) derived from *KRAS*-amplified biopsies maintained this genetic alteration in culture (**Figure 3E**, dark red). Despite their genetic stability, cells with inferred *KRAS* amplifications exhibited a profound phenotypic shift from basal *in vivo* to classical *ex vivo* (**Figure 3F**). Although selection of specific clones could play a role in this process, most of these models maintained high CNV similarity to their parent tumor at the early time point. For example, we observed that a CNV-defined clone from PANFR0575 with both *KRAS* and *GATA6* amplifications was plastic and shifted from strongly basal *in vivo* to classical in early organoid culture (**Figure 3G**). These observations provide strong evidence that phenotypic plasticity is an inherent feature of malignant PDAC cells and demonstrate that *KRAS* amplification alone is not sufficient to lock the basal state. Furthermore, they suggest that the tumor microenvironment can influence phenotype independent of genotype in this context.

### Transcriptional heterogeneity is shaped by the microenvironment

Given this strong phenotypic shift even for genetically similar samples, we next examined how *ex vivo* transcriptional phenotypes differed across our larger organoid cohort relative to their cognate patient samples. Globally, unbiased comparison of all malignant biopsy (7,078 cells) and organoid cells (n=14 models, 24,789 cells) revealed unique clusters for each sample and only two clusters that were admixed by donor. These admixed cells exhibited expression programs consistent with non-malignant stromal cells, had low overall CNV scores, and dissipated by later passages (**Supplemental Figure S5A-D**; **Methods**). Overall, samples with high tumor-averaged basal or IT phenotypes exhibited lower rates of long-term organoid propagation beyond passage 2 than models derived from classical tumors, where the majority established long-term cultures (**Figure 4A**). When comparing early passage CNV-confirmed organoids to their cognate patient tissue, culture in an *ex vivo* microenvironment caused greater deviation in transcriptional phenotype than CNV-defined genotype (**Figure 4A**, *P < 10^−6^* Student’s T test; **Methods**).

**Figure 4.**
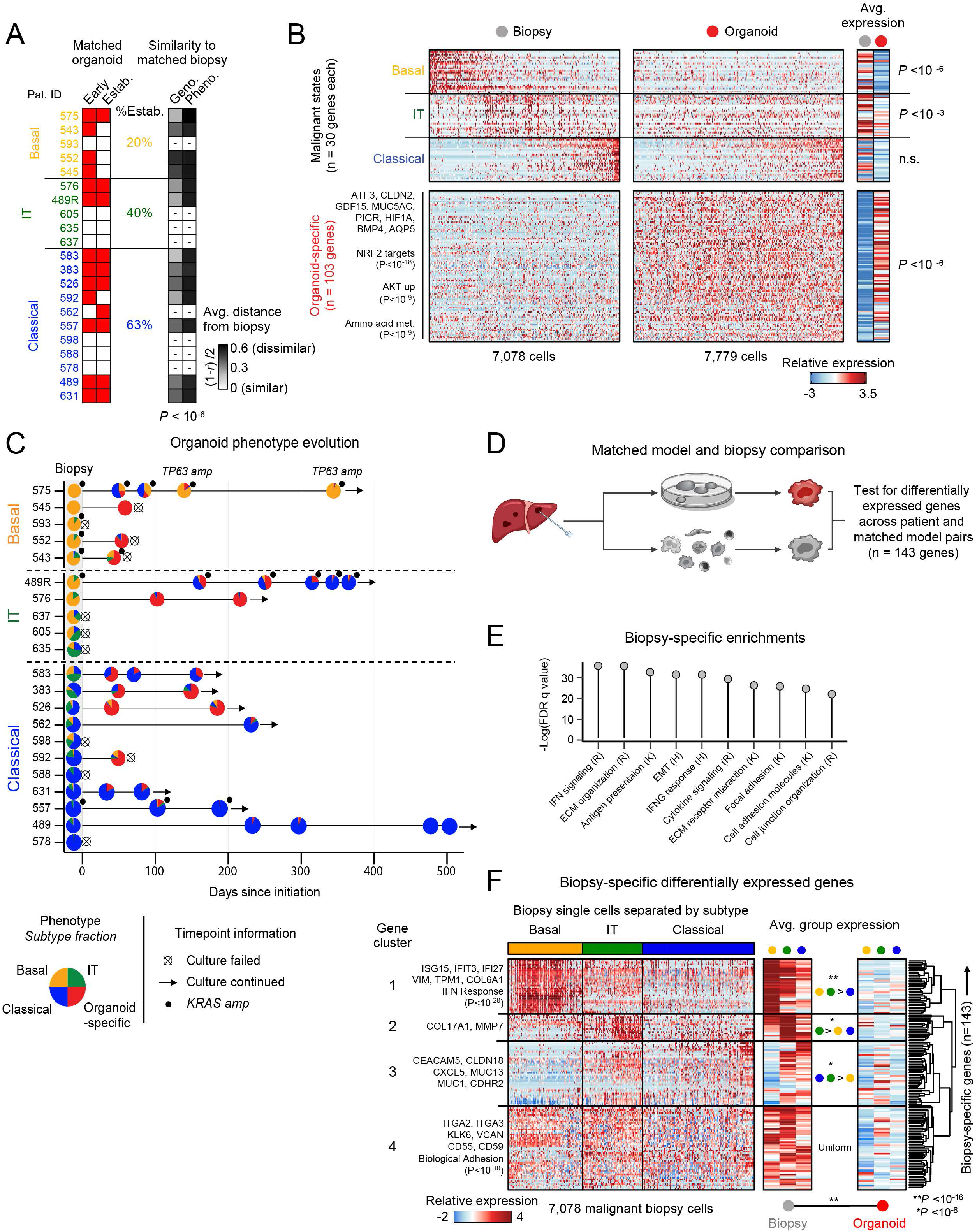
Organoid culture microenvironment selects against the basal state with phenotypic evolution over time. **(A)** Sampling from each model initiated as an organoid. Red fill represents measurements at an Early time point and if that biopsy established a long-term culture (Estab.). Right grey scale heat indicates the distance (**Methods**) between each biopsy-early organoid pair for CNVs (Geno.) or transcriptional subtype (Pheno.). *P*-value for Geno. vs Pheno. differences determined by student’s T test. **(B)** Relative expression for the malignant programs (top) and organoid-specific genes (bottom) in biopsy cells (left) and their matched, early passage organoid cells (n=13 models; right). Parenthetical *P*-values (left) indicate hypergeometric test for enrichment of pathways in the indicated gene clusters. Far right heat is average expression for all genes in each group, *P*-values determined by student’s T test. **(C)** Swimmer’s plot shows the evolution of organoid phenotype in the culture microenvironment. Each point indicates a passage when organoids were sampled with scRNA-seq, and pie chart fill indicates the fraction of single cells binned as each transcriptional subtype. **(D)** Schematic for matched tumor-organoid differential expression analysis. **(E)** Top differentially expressed genes *in vivo* (143 genes) are TME-associated and enrich for TME-associated pathways. All top enrichments shown are highly significant (*P*-value < 10^−12^). **(F)** Hierarchical clustering in biopsy cells (columns) of the relative expression for the 143 TME-associated genes preferentially expressed *in vivo* (rows). Cells are binned in the single-cell heatmap and the averages at right by their originating tumor’s average transcriptional subtype. Gene-level averages are split by biopsy (left) and organoid cells (right). Parenthetical *P*-values (left) indicate hypergeometric test for enrichment of pathways in the indicated gene clusters. For within-group differences in expression for biopsy averages, *P*-values are computed by one-way ANOVA followed by Tukey’s HSD and compare averaged expression of each gene cluster between cells from different biopsy subsets (middle heatmap; \**P*-value < 10^−8^; \*\**P*-value < 10^−16^). Overall biopsy versus organoid average expression difference for all 143 genes is determined by Student’s T test. *See also Supplemental Figure S5; Supplemental Table S5.*

We next assessed which specific tumor cell attributes contributed to phenotypic divergence in the *ex vivo* microenvironment. As with the *KRAS*-amplified samples, we observed a striking decrease in basal gene expression (*P < 0.000001*) and, to a lesser but still significant extent (*P < 0.001*), the IT program (**Figure 4B**, **top**). By contrast, aggregate classical gene expression remained largely unchanged in organoid conditions (**Figure 4B**, **top**). This loss of basal expression was surprising, given the more clinically aggressive and proliferative nature of basal tumors *in vivo* (**Supplemental Figure S3A**) (Connor et al., 2019; Moffitt et al., 2015). Organoid-specific gene expression features that were not present *in vivo* also emerged, including markers of epithelial identity, oxidative stress response pathways (e.g., NRF2 target genes), and amino acid metabolism (hereafter collectively referred to as “organoid-specific” gene expression; **Figure 4B**, **bottom**; **Supplemental Table S5**). In general, models assumed a more classical or organoid-specific phenotype over time in culture regardless of their parent tumor’s transcriptional identity (**Figure 4C**). Most models derived from basal or IT tumors exhibited early phenotypic deviation and cessation of growth within 100 days of initiation (e.g., PANFR0552; **Supplemental Figure S5E**) or outgrowth of only a sub-clone in culture (e.g., PANFR0489R; **Supplemental Figure S5F**). Classical tumors, meanwhile, tended to maintain their genotype and phenotype both early in culture and at later passages (e.g., PANFR0631; **Figure 4C**; **Supplemental Figure S5G**, clone A).

To better understand the contribution of clonal selection to this process, we performed linked genotype and phenotype assessment from iterative passages. We identified CNV-defined subclones in the parental biopsy and its associated serial organoid samples, and then assessed how the distribution of transcriptional states within each subclonal population evolved over time in culture (**Methods**). In both PANFR0489R and PANFR0575, *KRAS* amplification status remained invariant over time, but we observed significant phenotypic plasticity and clonal selection in both cases. In PANFR0489R, the predominantly basal clones *in vivo* rapidly decreased in abundance while other rarer clones with classical or organoid-specific phenotypes emerged as the dominant ones (**Supplemental Figure S5H**). In contrast, *in vivo* dominant clones from PANFR0575 were largely maintained at early passages but diverged significantly in their phenotype, transiently expressing more classical and organoid-specific phenotypes at passages 2 and 3 before eventually regaining basal transcriptional expression after >100 days in culture (**Supplemental Figure S5I**). Notably, the clones that came to dominate in PANFR0575 organoid culture (clones D and E, **Supplemental Figure S5I**) carried inferred *TP63* amplifications, a squamous-specifying transcription factor (Somerville et al., 2018), suggesting that certain genotypes, though rare, may still exert a strong effect despite opposing signals from the microenvironment. Taken together, these findings emphasize the importance of optimizing culture conditions and performing deep molecular characterization of patient-derived model systems to ensure faithful representation of the tumor.

Divergence from *in vivo* phenotype, despite relative similarity in genotype, suggested that the TME has a strong influence in determining PDAC cellular state. For each biopsy-organoid pair, we used differential expression to nominate transcriptional programs that were present *in vivo* but missing from *ex vivo* culture (**Figure 4D**; **Methods**). Broadly, genes preferentially expressed by malignant cells *in vivo* were related to soluble cytokine signaling, cell-cell communication, and tumor-microenvironment interactions, highlighting the absence of this crosstalk in organoid culture (**Figure 4E**). Hierarchical clustering revealed subtype-dependent expression patterns for these *in vivo*-specific genes (**Figure 4F**; **Supplemental Table S5**). For example, interferon response and EMT genes were significantly upregulated in basal and IT malignant cells *in vivo* (clusters 1 and 2, **Figure 4F**), while genes associated with cell-cell interactions and surface glycoproteins were more strongly expressed in IT and classical cells (cluster 3, **Figure 4F**). Genes related to biological adhesion were more uniform in their expression across the subtypes (cluster 4, **Figure 4F**). The relative absence of these TME-crosstalk genes in organoid culture and their differences in expression across transcriptional subtypes *in vivo* suggest that TME signals may play a role in specifying tumor cell phenotypes.

### Non-malignant composition of the metastatic microenvironment

The presence of TME-associated expression patterns in cancer cells *in vivo* suggested there may be subtype-dependent structure to, and instructive signaling from, the metastatic TME; however, relatively little is known about the structure and composition of the metastatic microenvironment in PDAC. We first analyzed the non-malignant cells (n=14,811) in the metastatic niche to further subclassify cell types and provide a more complete picture of the immune/stromal composition of metastatic disease (**Figure 5A**). Sub-clustering of T/NK cells revealed 4 cell types—*CD4*+ T, *CD8*+ T, NK, and *CD16+* (*FCGR3A+*) NK cells—each expressing the corresponding established markers (**Supplemental Figure S6A,B**; **Methods**). Similarly, an unsupervised examination within the monocyte/macrophage compartment revealed a tripartite continuum for tumor associated macrophages (TAMs), similar to one recently described in colorectal cancer, comprised of inflammatory *FCN1*+ “monocyte-like” TAMs, *C1QC*+ phagocytic TAMs, and *SPP1+* angiogenesis-associated TAMs (**Supplemental Figure S6C,D**; **Supplemental Table S2**) (Zhang et al., 2020; Zilionis et al., 2019). Representative marker expression across all previously described non-malignant cells is summarized in **Supplemental Figure S6E**.

**Figure 5.**
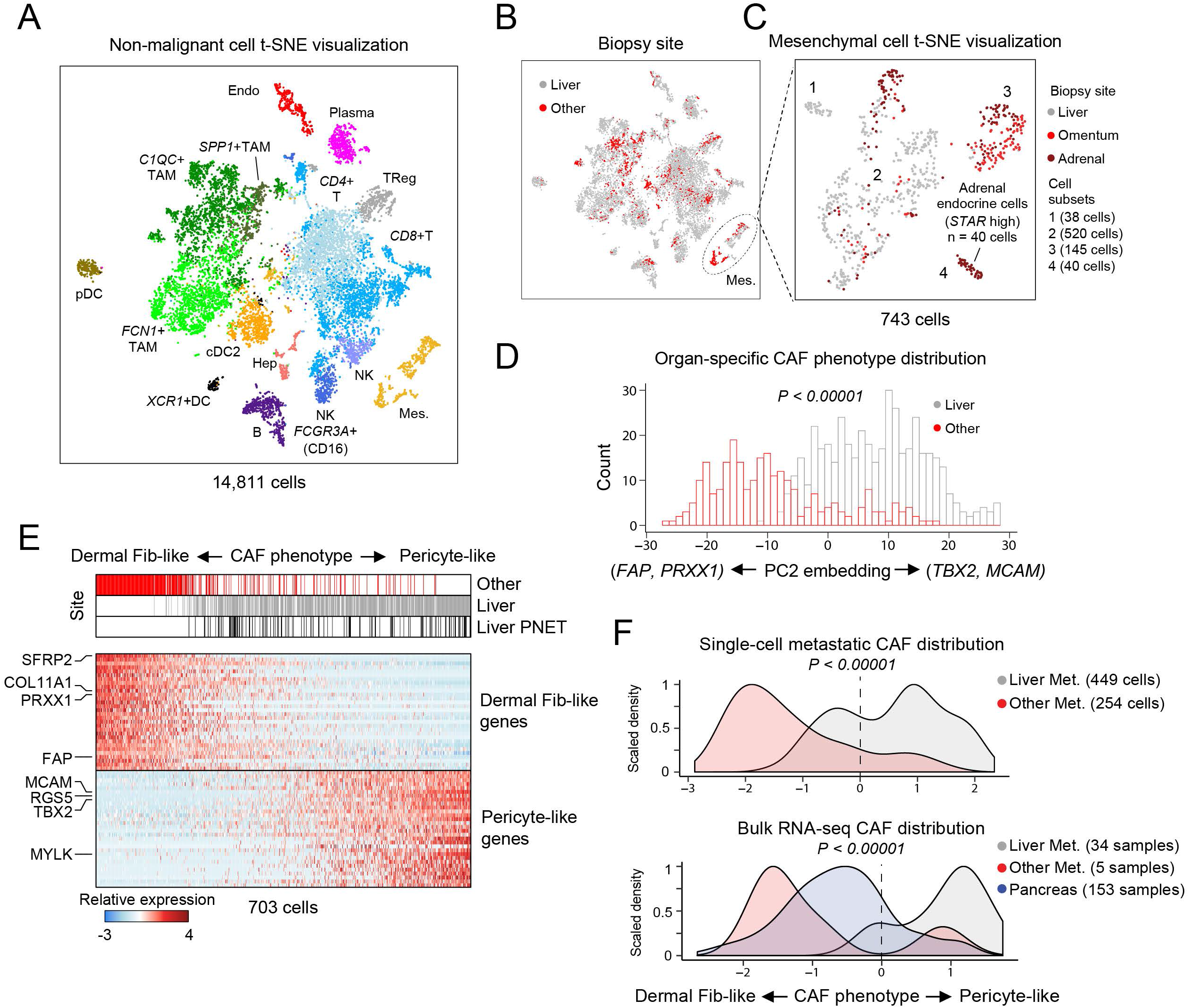
Immune heterogeneity and distinct fibroblast phenotypes exist in the liver metastatic microenvironment. **(A)** *t-*SNE visualization of non-malignant cells identified in the metastatic microenvironment, abbreviations are the same as in Figure 1D (TAM, tumor associated macrophage; NK, natural killer). **(B)** Same visualization as in **A**, but cells are colored by sampling site (Liver, grey; Other, red). Only the mesenchymal cells (dotted circle, Mes.) have appreciable separation by anatomical site. **(C)** *t-*SNE visualization of sub-clustering (SNN) performed on mesenchymal cells colored by their anatomical site. Cell subsets (1-4) determined by SNN clustering. **(D)** Frequency of CAFs (y axis, cell count) across PC2 scores, colored by site of biopsy tissue. *P*-value determined by student’s T test. **(E)** Heatmap for relative expression of the Dermal Fibroblast-like (PC2 low) and Pericyte-like (PC2 high) programs. Anatomical site is shown for each cell (top). **(F)** Density plots for CAF phenotype score in single cells from our metastatic cohort (top) or previously published PDAC bulk RNA-seq profiles (bottom) (Aguirre et al., 2018; Cancer Genome Atlas Research Network, 2017), fill indicates anatomical site. *P*-value determined by student’s T test (top) or by ANOVA followed by Tukey’s HSD (bottom). *See also Supplemental Figure S6; Supplemental Table S2.*

Although most samples in our cohort were taken from liver metastases (19/23), several originated from other sites including the omentum, adrenal gland, and peritoneum (**Figure 1B**, **“**other”). Interestingly, while we found equal distribution of immune cells among the anatomical sites, mesenchymal cell populations clustered predominantly by the location of the metastatic lesion (**Figure 5B,C**). Excluding adrenal-specific endocrine cells (**Figure 4C**; subset 4, 40 cells), we identified 3 mesenchymal subclusters with relatively uniform expression for canonical cancer-associated fibroblast (CAF) markers (**Figure 5C**; **Supplemental Figure S6F**). PCA of these cells revealed a continuum of states along PC2, with uniform expression of the previously described myofibroblast (myCAF) signature (Elyada et al., 2019; Ohlund et al., 2017) but further separating into cells favoring high expression of dermal fibroblast-like genes (PC2 low, *FAP, PRXX1, SFRP2*) or pericyte-like genes (PC2 high, *RGS5, MCAM, TBX2*; **Supplemental Figure S6G-I**; **Supplemental Table S2**) (Ascension et al., 2020; Bartoschek et al., 2018; Di Carlo and Peduto, 2018; Hosaka et al., 2016; Pelon et al., 2020; Philippeos et al., 2018). PC3 described a small subset of cells, derived largely from a single tumor (PANFR0489R), that were highly consistent with the previously established inflammatory fibroblast (iCAF) program (Elyada et al., 2019; Ohlund et al., 2017) (**Supplemental Figure S6H-J**).

While tumors from each location contained both mesenchymal subsets, we noted a strong organ-specific skewing along PC2 with pericyte-like phenotypes being preferentially associated with liver biopsies (**Figure 5D,E**; **Supplemental Figure S6K**). To validate these observations in larger cohorts, we assessed bulk RNA-seq datasets using these dermal fibroblast- and pericyte-like CAF signature scores and observed a similar predilection for the pericyte-like expression program in liver metastases (**Figure 5F**; **Methods**). Interestingly, tumors in the pancreas (n = 153 samples) favored expression of the dermal fibroblast-like program, suggesting a substantially different mesenchymal microenvironment in primary versus liver metastatic PDAC (**Figure 5F**). Thus, we observed diverse immune and stromal cell types in the metastatic TME and identified site specific mesenchymal features unique to the liver metastatic niche compared with primary disease.

### Transcriptional subtypes associate with distinct immune microenvironments

After cataloging the cell types in the metastatic TME, we searched for associations between malignant subtype and the immune microenvironment. For each tumor sample, we first computed the fractional representation of each non-malignant cell type per biopsy. Five tumors were excluded from this analysis on the basis of low cell counts (<200 cells) or indeterminant transcriptional subtype (PanNET or no tumor cells captured; **Supplemental Figure S6L**). To describe the overall microenvironmental composition for each tumor, we applied Simpson’s diversity index, a measure of biodiversity commonly used in ecology to describe the number of species (cell types) present in an ecosystem (tumor) and their relative abundance. We observed that tumors with more classical or IT phenotypes exhibited greater microenvironmental diversity, while strongly basal tumors had a more homogeneous TME (**Figure 6A**). Hierarchical clustering over the relative abundance of each non-malignant subset across the biopsy cohort revealed the specific cell types driving these overall diversity differences (**Figure 6B,C**). Specifically, C1QC+ TAMs dominated the microenvironments of strongly basal tumors, and both CD8+ and CD4+ T cells were significantly depleted in basal contexts compared to the rest of the samples in the cohort (**Figure 6C,D**). T cells most often originated from biopsies with higher IT malignant fractions (**Figure 6B,C**) and their abundance was positively correlated with this malignant phenotype in our cohort (**Figure 6E**). We also broadly observed these patterns within TCGA bulk RNA-sequencing data of other epithelial malignancies (Cancer Genome Atlas Research et al., 2013), where we observed evidence for reduced levels of immune-related gene expression in tumors with high basal/squamous gene expression (**Supplemental Figure S6M**, cluster 4). Taken together, these findings suggest coordinated interactions between malignant phenotypes and the local TME with decreased immune cell diversity and a greater degree of immune exclusion associated with basal contexts (**Figure 6F**).

**Figure 6.**
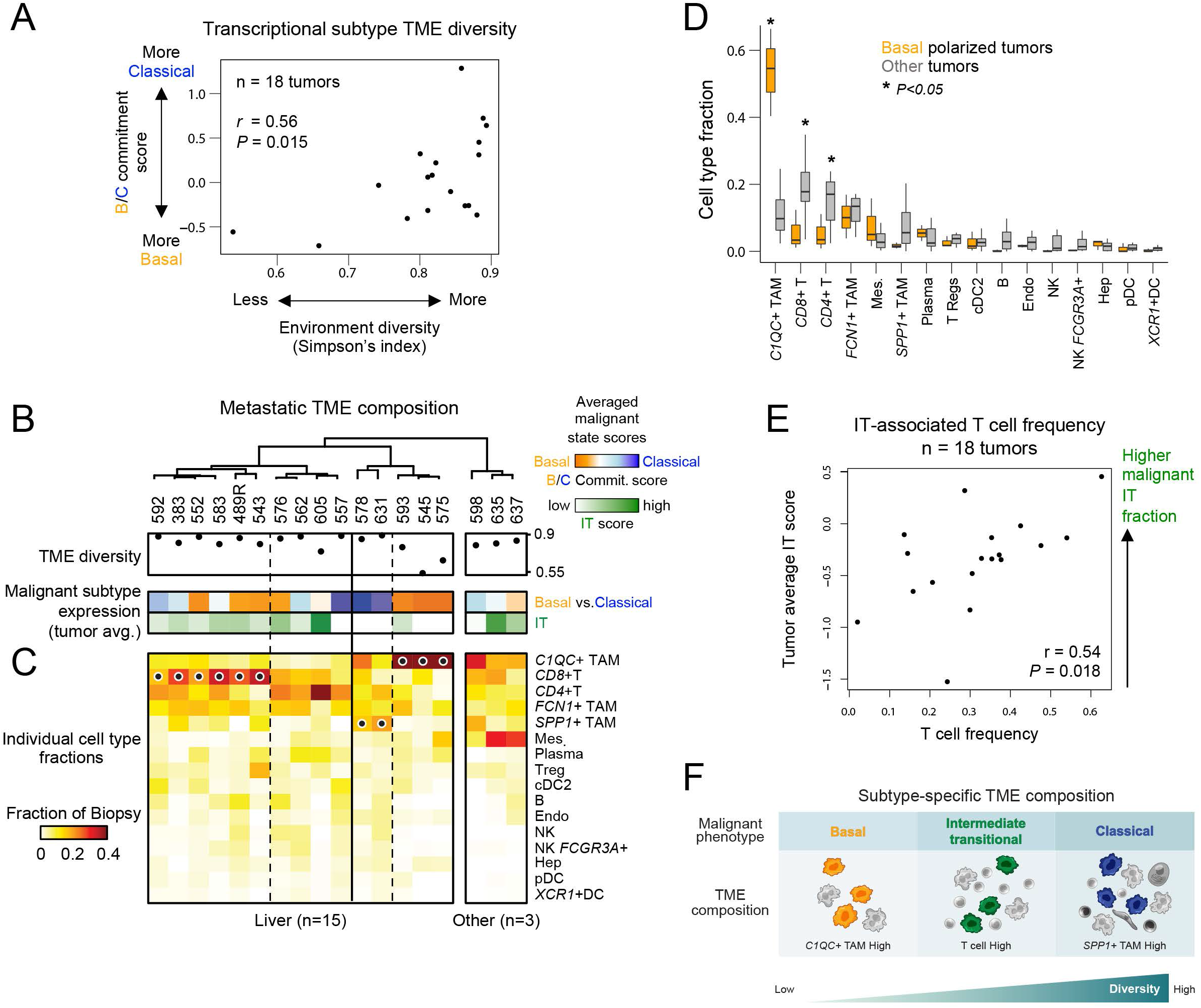
Transcriptional subtypes associate with distinct metastatic microenvironments. **(A)** Correlation between microenvironment diversity (Simpson’s Index, x axis) and the average malignant basal-classical commitment score for each biopsy (y axis). **(B)** Dot plot indicates the Simpson’s Index calculated for each biopsy and heat bars indicate each tumor’s average malignant cell expression for each of the malignant transcriptional programs. **(C)** Fraction of each non-malignant cell type (heat, rows) in each biopsy sample (columns). Dots indicate top statistically significant cell type frequency differences calculated using Kruskal-Wallis test with multiple hypothesis correction. Samples are ordered as in **B.** **(D)** Box plots compare cell type fraction between the basal polarized tumors with low diversity (PANFR0593, 575, 545) and all others. *P*-value determined by student’s T test. **(E)** Correlation between T cell fraction and IT malignant score. **(F)** Schematic summarizing associations between microenvironmental diversity, non-malignant infiltrates, and tumor subtype. *See also Supplemental Figure S6; Supplemental Table S2.*

### The soluble microenvironment shapes PDAC cellular phenotypes

Based on our observations that: 1) the microenvironment influences malignant phenotype independent of genotype; 2) gene expression programs associated with cytokine signaling, EMT, and cell-cell interaction are enriched *in vivo* but missing from cells cultured as organoids; and, 3) malignant states and immune cell infiltration are coordinated in a subtype-specific manner, we hypothesized that incorporation of soluble factors specific to the TME of each transcriptional subtype may drive tumor cell state shifts (**Figure 7A**). Complete PDAC organoid media (**Supplemental Table S6**) (Boj et al., 2015; Tiriac et al., 2018) contains various growth factors that could skew malignant transcriptional state, so we first tested the effects of withdrawing various soluble factors. We cultured four organoid models in media without any additives (“Minimal” media, containing only Glutamax, anti-microbials, HEPES buffer, and Advanced DMEM/F12 media; **Figure 7B**; **Supplemental Table S6**; **Methods**). We observed a robust increase in basal gene expression and a decrease in organoid-specific gene expression in specimens cultured for 6 days in minimal media relative to those in complete organoid media (“Complete”, **Figure 7B**). Although we found that the fraction of cycling cells in minimal media decreased, the organoids continued to grow under these conditions and exhibited stable CNV profiles, indicating that these responses were unlikely to be driven by acute selection (**Supplemental Figure S7A,B**). We cultured one model, PANFR0562, in minimal media for a longer duration and observed that the phenotypic distribution shifted even further toward IT and basal phenotypes (**Figure 7C**), demonstrating that recovery of all three states is possible *ex vivo*. Since minimal medium lacks both serum and mitogens to support prolonged cell growth, we also tested whether culturing organoids in a reduced organoid media formulation (“OWRNA”, complete organoid media with removal of WNT3A, RSPONDIN-1, NOGGIN, and A83-01; **Supplemental Table S6**; **Methods**) supported proliferation while allowing expression of basal and IT phenotypes. We found that organoids maintained under OWRNA conditions began to express basal and IT features while also strengthening classical gene expression and continuing to proliferate (**Supplemental Figure S7C**).

**Figure 7.**
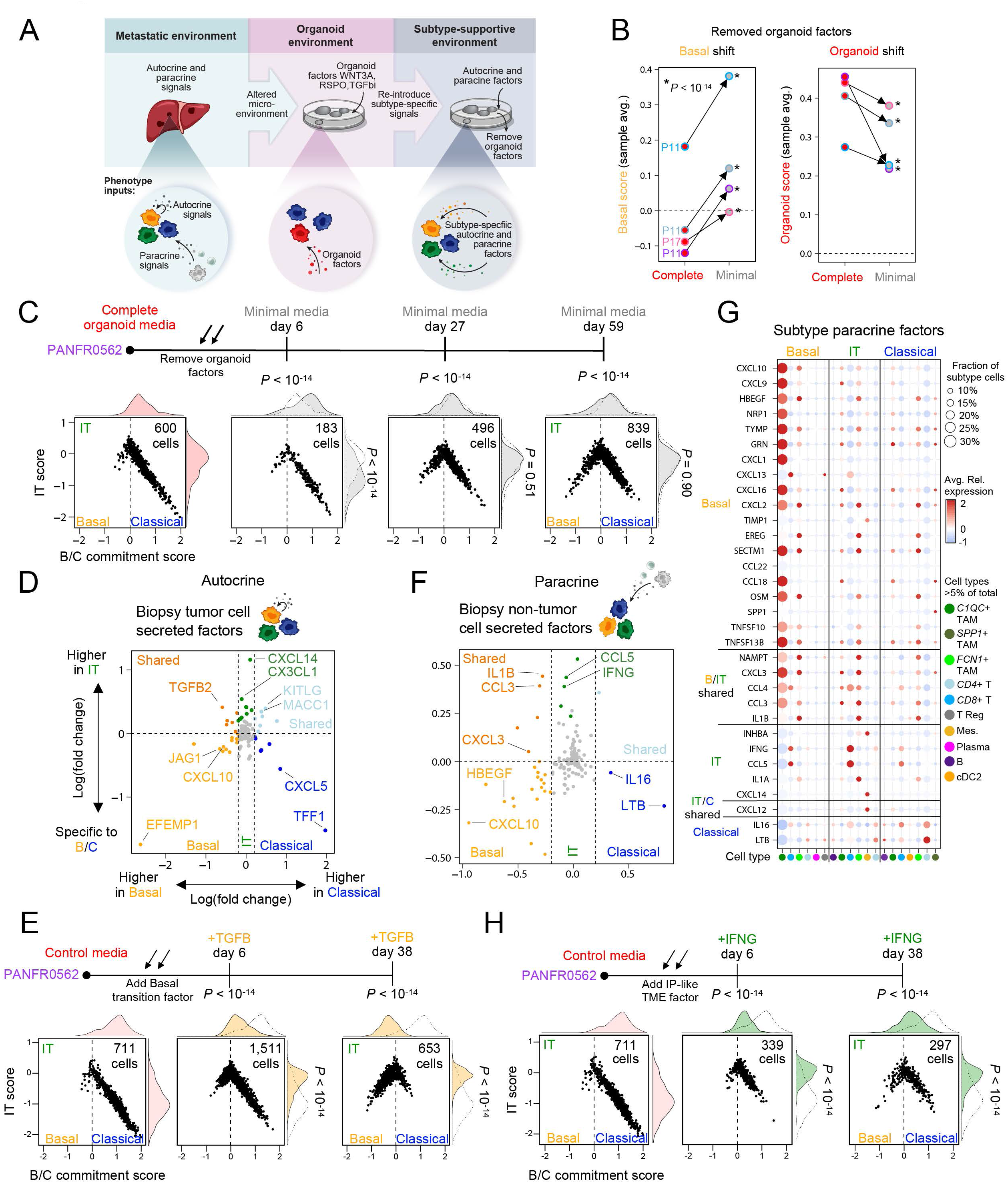
Tumor subtype-specific secreted microenvironmental factors rescue malignant transcriptional heterogeneity. **(A)** Schematic describing microenvironmental inputs *in vivo* (“Metastatic environment”) versus *ex vivo* (“Organoid environment”) to tumor phenotype. Right panel (“Subtype-supportive environment”) describes an overall strategy to recover malignant transcriptional heterogeneity by removing organoid factors (**B, C**) and adding state-specific autocrine (**D, E**) or paracrine (**F-H**) factors. **(C)** Tied dot plot represents the sample average basal score (left) and organoid-specific score (right) in the indicated conditions. Lines tie samples and color outlines indicate sample identity as in Figure 1B. *P*-value compares respective single cell distributions within models and was calculated by student’s T test. **(D)** Cell state diagrams for organoid cells cultured in complete medium or at 3 time points in minimal media. *P*-values for group differences between B/C commitment (top) and IT scores (right) were calculated by ANOVA followed by Tukey’s HSD. *P*-values displayed are for that timepoint vs. the complete media condition. **(E)** Differential expression (Wilcoxon rank sum test) for known secreted factors by *in vivo* tumor cells (autocrine) between basal and classical (x axis) and IT malignant cells and the rest (y axis). Subtype-specific genes that pass significance after multiple hypothesis correction (*P < 0.05*) are colored by their group association. **(F)** Cell state diagrams with marginal density plots for organoid cells cultured in control medium (OWRNA, reduced organoid medium) or at 2 time points in control media with TGF-β. *P*-values for group differences between B/C commitment (top) and IT scores (right) were calculated by ANOVA followed by Tukey’s HSD. *P*-values displayed are for that timepoint vs. the control media condition. **(G)** Differential expression (Wilcoxon rank sum test) for known secreted factors by all non-malignant cells (paracrine) found in basal and classical (x axis) and IT biopsies and the rest (y axis). Subtype-specific genes expressed by non-malignant cells that pass significance after multiple hypothesis correction (*P < 0.05*) are colored by their group association. **(H)** Dot plot for the subtype-specific significant differentially expressed paracrine factors. Subtype-specific non-malignant cell types (columns) and significant genes (rows) are binned by subtype association as in Figure 6C and Figure 7F. Dot size represents that cell type’s fraction within tumors of each subtype, and fill color indicates average expression. Only cell types with a fractional representation >5% from each subtype are visualized. **(I)** Cell state diagrams with marginal density plots for organoid cells cultured in control medium (OWRNA, reduced organoid medium, as in **E**) or at 2 time points in control media with IFNγ. *P*-values for group differences between B/C commitment (top) and IT scores (right) were calculated by ANOVA followed by Tukey’s HSD. *P*-values displayed are for that timepoint vs. the control media condition. *See also Supplemental Figure S7; Supplemental Table S6 & S7.*

To assess whether these microenvironment-driven effects on transcriptional states were specific to organoid models or also observed in other cell culture models, we examined PDAC cell lines, as these are also commonly used to study PDAC biology but are grown in different culture conditions. We compared bulk RNA expression data from patient tumors (n=219) (Aguirre et al., 2018; Cancer Genome Atlas Research Network, 2017), our own organoid cohort (n=44), and established cell lines (n=49, CCLE) (Barretina et al., 2012; Ghandi et al., 2019) and observed strong culture method-dependent phenotypic skews wherein most organoid models expressed classical phenotypes while cell lines exhibited basal phenotypes (**Supplemental Figure S7D,E**). This observation suggests neither platform accurately represents the full repertoire of transcriptional states seen in patients and provides additional evidence that environmental conditions can profoundly influence transcriptional state. We ruled out the effects of extracellular matrix dimensionality from media formulation by culturing established 3-dimensional (3D) organoid models as 2-dimensional (2D) cell lines on tissue culture plastic in the same organoid media— this had little effect on transcriptional subtype across the models tested (**Supplemental Figure S7F**). Next, we took each model type (cell lines and organoids) and cultured it in the reciprocal media condition to ask whether media alone could influence transcriptional subtype. Organoid cells grown in standard cancer cell line medium (“RP10”, RPMI-1640 with 10% fetal bovine serum) gained expression of basal programs (**Supplemental Figure S7C**), while CFPAC1 (an established PDAC cell line) lost basal and classical features and gained organoid-specific gene expression when grown in complete organoid media (“Complete media”, **>Supplemental Figure S7G**). Taken together, these findings demonstrate that the microenvironment is an instrumental contributor to shaping malignant phenotypes in PDAC. Moreover, the cell state plasticity suggests the possibility of testing subtype-specific conditions to support the full repertoire of *in vivo* phenotypes.

### Applying subtype-specific TME signals drives patient-relevant subtype heterogeneity

Finally, we hypothesized that specific factors from subtype-specific TMEs could recover clinically relevant transcriptional heterogeneity *ex vivo* (**Figure 7A**). *In vivo,* the secreted factor milieu surrounding tumor cells originates from at least two sources that may influence malignant phenotype: tumor cells themselves (“autocrine” factors) and non-tumor cells (“paracrine” factors, **Figure 7A**). First, to nominate possible autocrine signals, we identified tumor cell secreted factors specific to the three subtypes and noted distinct cytokines expressed by each (**Figure 7D**; **Supplemental Table S7**; **Methods**). Since malignant cells derived from predominantly basal and IT tumors lose their phenotype in organoid culture, we first tested factors specific to IT and basal states *in vivo*. *TGFB2* was the top differentially expressed secreted factor shared by tumor cells in both basal and IT TMEs (**Figure 7D**). Organoids cultured with TGF-β ligands exhibited a loss of classical expression programs and a near complete shift toward IT and basal phenotypes, matching what we observed *in vivo* (**Figure 7E**). Reemergence of basal phenotypes in both minimal media (**Figure 7C**), and TGF-β conditions (**Figure 7E**) suggest that different types of microenvironmental pressure can lead to the basal phenotype. Moreover, they suggest that culture conditions can be tuned to achieve compositional differences spanning pure classical, heterogenous, and pure basal phenotypes, akin to those seen *in vivo*.

Using a similar approach, we next searched for differentially expressed paracrine factors supplied by the non-tumor cells in the TME from each subtype. Here, we noted an increasing number of differentially expressed factors in IT and basal contexts, likely reflecting the specific immune cell type enrichments: TAM and T cell dominant in basal and IT TMEs, respectively (**Figure 6A-C**; **Figure 7F**; **Supplemental Table S7**). We then mapped each subtype-specific paracrine factor to its cognate cell type to summarize the overall cell type and secreted factor combinations that shape the subtype-specific TMEs in metastatic PDAC (**Figure 7G**). Interestingly, we found that *IFNG* originating from *CD8+* T cells was most highly expressed in the IT TME (**Figure 7F,G**). This was consistent with a relatively higher T cell fraction in IT tumors (**Figure 6B,F**) and the relative increase in IFN responsive gene expression in IT and basal tumor cells (**Figure 4E,F**). Given these corroborating correlative data, we directly tested whether exogenous IFNγ could induce transcriptional plasticity towards an IT state. Cells exposed to IFNγ showed a dramatic shift toward the IT state with concomitant decrease in expression of classical signatures (**Figure 7H**). In contrast with exogenous TGF-β (**Figure 7E**), microenvironmental IFNγ seemed to more specifically induce an IT state, as these cells did not fully transition to basal phenotypes at later timepoints (**Figure 7H**). These findings demonstrate that the microenvironment plays a critical role in specifying tumor transcriptional phenotypes and provide evidence for significant PDAC tumor cell plasticity in response to microenvironmental cues.

## DISCUSSION

Here, by linking single-cell profiling of *in vivo* patient specimens to matched organoid models, we have built an essential comparative dataset to disentangle the contributions of cell-intrinsic versus -extrinsic factors to cancer cell transcriptional states in metastatic PDAC. We leveraged the precision afforded by scRNA-seq to identify a new PDAC cell state that co-expresses the basal and classical programs and behaves as a transitional intermediate between the basal and classical subtypes. Importantly, the identification of large fractions of co-expressing IT cells in human tumor biopsies using both mIF and scRNA-seq suggests interconversion between the classical and basal subtypes occurs frequently in response to various cues *in vivo* and implies that this intermediate state may be a hallmark of intratumoral plasticity and tumor cell transcriptional evolution. In fact, in contrast to prior reports (Chan-Seng-Yue et al., 2020), all tumors that had mixed but discrete populations of basal and classical cells also exhibited proportional fractions of co-expressing IT cells. Our matched organoid studies provide strong evidence that this extensive transcriptional heterogeneity is heavily influenced by the microenvironment, a finding that is further reinforced by the identification of subtype-dependent TME structure. As such, this work provides a detailed description of the PDAC metastatic niche, critical insight into the role of the microenvironment in determining cancer cell phenotype in PDAC, and a general framework for discovering and manipulating these relationships across cancer contexts.

Although mutations in *KRAS* play a critical role in pancreatic oncogenesis, PDAC cells have also been shown to adopt more RAS-independent phenotypes as a mechanism of resistance to KRAS suppression (Muzumdar et al., 2017). Our findings help to reconcile these opposing observations by suggesting that KRAS target gene expression is more strongly associated with the IT cell state than either basal or classical extremes. This finding suggests that while upregulation of KRAS signaling by amplification or other mechanisms may play an important role in the transition toward the basal state (Chan-Seng-Yue et al., 2020; Miyabayashi et al., 2020), it may become less functionally important once this state transition is complete. Furthermore, the presence of IT cells enriched for *KRAS* and inflammatory response gene expression is reminiscent of phenotypes seen in mouse models that suggest inflamed progenitor-like cells as those that tolerate *KRAS* mutations and initiate tumorigenesis (Alonso-Curbelo et al., 2021; Li et al., 2021).

Our single-cell data support the association between *KRAS* amplifications and the basal state *in vivo*; however, when we compared our matched *KRAS* amplified biopsy and organoid cells, we saw that this genotype did not lock cells into the basal state, and that microenvironmental conditions were a dominant factor in determining tumor cell transcriptional subtype. Serial sampling of organoid models across successive passages demonstrated both phenotypic drift and sub-clonal outgrowth, mirroring the genetic evolution of PDXs and cell lines in culture (Ben-David et al., 2017; Ben-David et al., 2018), and highlighting the complex interplay between genetics and microenvironmental influences on transcriptional plasticity and clonal selection. This facile transition between subtypes has important implications for drug treatment, and future studies using lineage tracing approaches are needed to better understand the evolutionary dynamics in this system and how to track and exploit these processes therapeutically. Additional studies into the epigenetic regulatory mechanisms underlying PDAC state transitions will also be a critical next step in further delineating the relationships between genotype, microenvironment, and phenotype.

Although we have identified co-expressing IT cells in both primary and metastatic tumors, the transcriptional programs associated with co-expression may differ between these contexts. We hypothesize that the basal state may be a common phenotypic endpoint for PDAC tumor cells in response to microenvironmental stress, with superimposed transcriptional variation depending upon the specific stressors a given tumor cell must overcome to reach this state. Supporting this concept is the observation that cells exhibiting basal phenotypes show concomitant expression of EMT, IFN response, or hypoxia response signatures, and these expression patterns may be driven by the specific microenvironment (Benci et al., 2016; Connor et al., 2019). In addition, our finding that diverse microenvironmental signals, including nutrient deprivation (“Minimal media”), autocrine and stromal signals (TGF-β), and immune signals (IFNγ), induce the transition away from the classical subtype further supports this conclusion. We postulate that IT intermediates likely house similar context-dependent complexity depending on the tissue of residence.

Similar to malignant cells, the non-malignant cell types in the metastatic TME were varied in phenotype and overall composition. Although we mainly sampled liver metastases, we identified strong differences between mesenchymal populations from different biopsy sites. We observed that the liver metastatic niche was enriched for pericyte-like myofibroblasts (Bartoschek et al., 2018; Di Carlo and Peduto, 2018; Hosaka et al., 2016; Pelon et al., 2020), while other sites of metastasis and primary disease were enriched for dermal fibroblast-like phenotypes. Given the pivotal role that has been suggested for the fibrotic TME in primary disease (Ho et al., 2020; Sahai et al., 2020), these findings carry important implications for targeting the stromal compartment in primary versus metastatic PDAC. For example, inhibitors targeting FAP have recently shown preclinical efficacy (Fabre et al., 2020), but we observe FAP expression favors dermal fibroblast-like cells but not pericyte-like myofibroblasts which are more prevalent in liver metastases. As such, examination of these fibroblast phenotypes across larger sample sets may help to identify additional clinically relevant variation in tumor-fibroblast crosstalk, and site-specific combinatorial strategies may be needed to effectively target the PDAC tumor stroma.

We show how scRNA-seq can be employed to define the structure of the metastatic niche and uncover formerly unappreciated relationships between tumor transcriptional phenotype and the local TME. Although traditionally thought of as a uniformly “immune-cold” tumor, our findings highlight that the immune microenvironment in metastatic PDAC harbors a layer of complexity closely linked to tumor cell transcriptional subtype that may provide new avenues for therapeutic targeting. Notably, we observed high levels of *IFNG* expression by CD8+ T cells and coordinated elevation in IFN response gene expression in IT and basal malignant cells. We recapitulated this shift from a classical to a more IT state in organoid models exposed to IFNγ, suggesting that malignant adaptation to signals from the TME may contribute to driving IT and basal phenotypes. Similar to the relationship between inflammation and tumorigenesis (Alonso-Curbelo et al., 2021; Li et al., 2021), we speculate that as tumors become inflamed and immune-activated, malignant cells display enhanced plasticity, transition to an IT state in response, and then progress to a fully basal phenotype with concomitant immune evasion and exclusion. These relationships may have implications for PDAC response to immunotherapy given that a productive immune response may promote more aggressive basal phenotypes (Benci et al., 2016; Li et al., 2018). Notably, we observed evidence for basal expression signatures with a corresponding paucity of immune cell type signatures in multiple other epithelial cancers, suggesting that coordination of malignant and immune responses in basal contexts may be a broadly relevant phenomenon across many cancer types. Additional studies with co-culture, mouse models, or serial samples from patients on active immunotherapy may further clarify these coordinated and reciprocal tumor-immune interactions.

More generally, our approach using matched *in vivo* malignant populations as a reference for *ex vivo* perturbations and model generation provides a critical framework for understanding the signals that drive clinically relevant phenotypes but are missing from organoid and cell line cultures. The genetic evolution of *ex vivo* models is a well-established phenomenon which carries functional consequences (Ben-David et al., 2019; Ben-David et al., 2018). Our study highlights similar *ex vivo* evolution for transcriptional variation, but also provides a strategy to rescue malignant phenotypes by re-introduction of soluble signals needed for their support *in vivo*. This approach may offer a more tractable system for state-specific high throughput screening compared with more complex heterotypic co-cultures or PDX systems. With a catalogue of matched *in vivo* phenotypes as a reference, this workflow empowers not only model fidelity, but enhances our ability to learn the phenotypic boundary conditions for individual tumors. For example, we can begin to define whether certain pressures induce cell state transitions in specific subsets of *ex vivo* models and identify which combinations of factors impede or synergistically enhance these transitions. Furthermore, these studies highlight how model generation in different growth contexts—organoids, cell lines, spheroids—may lead to the identification of emergent tumor cell properties. Learning these rules across different tumor contexts and understanding which non-malignant cell types participate *in vivo* would allow for the full appreciation of the symbiotic relationships within tumor ecosystems and provide a valuable foundation for leveraging microenvironmental manipulation to control tumor cell phenotype and behavior.

In sum, our data demonstrate coordinated phenotypic evolution driven by reciprocal interactions between malignant cells and the TME in PDAC. Just as we consider therapeutic combinations to target tumor cell intrinsic properties, paracrine interactions with the TME may equally drive tumor cell phenotype and thus require consideration in designing combination strategies. We provide a framework for relating malignant cells, the TME, and patient-derived model systems that may be applicable in other tumor types with clinically relevant transcriptional variation across the malignant and microenvironmental compartments.

## ACKNOWLEDGEMENTS

We thank the study participants and their families for enabling this research. This work was funded by the Lustgarten Foundation Dedicated Laboratory Program (B.M.W., A.J.A.), Dana-Farber Cancer Institute Hale Center for Pancreatic Cancer Research (A.J.A., B.M.W., W.C.H.), the Doris Duke Charitable Foundation (A.J.A.), Pancreatic Cancer Action Network (A.J.A., B.M.W.), NIH-NCI K08 CA218420-02 (A.J.A.), P50 CA127003 (A.J.A., B.M.W.), U01 CA176058 (W.C.H.), U01 CA224146 (W.C.H., A.J.A.), U01 CA210171 (B.M.W.), U01 28020510 (W.C.H., A.K.S.), 1U2CCA23319501 (A.K.S.), U54 CA217377 (W.C.H., S.R.M., A.K.S.), U01 CA250549 (W.C.H., S.R., A.J.A.), Stand Up To Cancer (B.M.W.), Noble Effort Fund (B.M.W.), Wexler Family Fund (B.M.W.), Promises for Purple (B.M.W.), Hope Funds for Cancer Research Postdoctoral Fellowship (S.R.), Harvard Catalyst/The Harvard Clinical and Translational Science Center UL1 TR002541 (S.R.), Koch Institute for Integrative Cancer Research at MIT and the Dana-Farber/Harvard Cancer Center Bridge Project (W.C.H., S.R.M., S.R.), Dana-Farber Cancer Institute Gloria Spivak Faculty Advancement Fund (S.R.), Hopper-Belmont Foundation Inspiration Award (S.R.), Marcotte Center for Cancer Research (B.E.J.), Finnish Cultural Foundation (S.A.V.), Orion Research Foundation sr (S.A.V.), Broman Family Fund for Pancreatic Cancer Research (K.N.), Ludwig Center for Molecular Oncology at MIT (A.W.N.), Searle Scholars Program (A.K.S.), Beckman Young Investigator Program (A.K.S.), Sloan Fellowship in Chemistry (A.K.S.), and the Pew-Stewart Scholars Program for Cancer Research (A.K.S.). We also thank Bryan Bryson for helpful discussions of macrophage phenotypes.

## AUTHOR CONTRIBUTIONS

S.R., P.S.W., B.M.W., W.C.H., A.J.A., and A.K.S. conceived and led the study. S.R., P.S.W., A.W.N., H.L.W., R.L.K., J.G.R., K.E.L., N.M., M.S.R., R.W.S.N., J.W., I.W., A.R., and E.S. designed and conducted experiments. S.R., P.S.W., A.W.N., H.L.W., A.D., A.A.B., K.S.K., A.M.J., L.F.S., Y.Y.L., A.D.C., and J.M.M. analyzed and interpreted data. S.A.V., A.D.C., E.R., D.R., L.K.B., A.R., A.M.J., T.E.C., K.P., D.A.R., K.N., J.M.C., and J.A.N. contributed patient samples, clinical data, and intellectual input. B.E.J., T.J., L.C., J.A.N., S.R.M., B.M.W., W.C.H., A.J.A., and A.K.S. supervised aspects of the project. S.R. and P.S.W. wrote the manuscript with input from A.W.N., H.L.W., S.R.M., B.M.W., W.C.H., A.J.A., and A.K.S. and review by all other co-authors.

## METHODS

### RESOURCE AVAILABILITY

#### Lead Contact

Further information and requests for resources and reagents should be sent to and will be fulfilled by Dr. Alex Shalek.

#### Data Availability

The single-cell RNA sequencing data reported in this paper will be deposited in a central data sharing repository (Genomic Data Commons) under the NCBI Database of Genotypes and Phenotypes (dbGaP). Code will be available upon request.

##### Tissue collection and dissociation

Investigators obtained written, informed consent from patients with pancreatic cancer for Dana-Farber/Harvard Cancer Center Institutional Review Board (IRB)-approved protocols 11-104, 17-000, 03-189, and/or 14-408 for tissue collection, molecular analysis, and organoid generation. Core needle biopsy specimens were collected and the first core was sent for pathologic analysis. One or more additional cores were then allocated for scRNA-seq and organoid generation.

Samples were minced into small portions using a scalpel and then digested at 37°C for 15 minutes using digest medium that consisted of human complete organoid medium (see below), 1 mg/mL collagenase XI (Sigma Aldrich), 10 μg/mL DNase (Stem Cell Technologies), and 10 μM Y27632 (Selleck) (Tiriac et al., 2018). In our initial process optimization, we found that dissociation times below 30 minutes, while not always completely digesting all biopsy material and potentially affecting the representation of difficult to dissociate cell types (e.g., fibroblasts), resulted in greater cell viability and improved RNA quality downstream. After digestion, cells were washed, treated with ACK lysing buffer (Gibco) to lyse red blood cells, washed again, and counted using a hemocytometer with 0.4% Trypan blue (Gibco) added at 1:1 dilution for viability assessment. We allowed residual tissue chunks to settle before selecting a predominance of single cells for counting and Seq-Well processing. We allocated between 10,000 and 15,000 viable cells per Seq-Well array based upon total cell counts, and where possible we prepared two arrays per sample. Most samples were processed and loaded onto Seq-Well arrays within 2-3 hours of biopsy acquisition.

##### Organoid generation and sampling

Cells remaining after scRNA-seq allocation were initiated and maintained as patient-derived organoid cultures as previously described (Boj et al., 2015; Tiriac et al., 2018). In brief, digested cells were seeded in 3-dimensional (3D) Growth-factor Reduced Matrigel (Corning) and fed with human complete organoid medium containing advanced DMEM/F12 (Gibco), 10 mM HEPES (Gibco), 1x GlutaMAX (Gibco), 500 nM A83-01 (Tocris), 50 ng/mL mEGF (Peprotech), 100 ng/mL mNoggin (Peprotech), 100 ng/mL hFGF10 (Peprotech), 10 nM hGastrin I (Sigma), 1.25 mM N-acetylcysteine (Sigma), 10 mM Nicotinamide (Sigma), 1x B27 supplement (Gibco), R-spondin1 conditioned media 10% final, Wnt3A conditioned media 50% final, 100 U/mL penicillin/streptomycin (Gibco), and 1x Primocin (Invivogen) (**Supplemental Table S6**). 10 μM Y27632 (Selleck) was included in the culture medium of newly initiated samples until the first media exchange. For propagation, organoids were dissociated with TrypLE (Gibco) before re-seeding into fresh Matrigel and culture medium.

After initial processing of fresh tissue specimens, we monitored samples closely for organoid growth. We did not passage organoids at set time intervals, as there was significant variability in the time needed to establish relatively robust growth of organoids (**Figure 4C**). Instead, we maintained early passage organoids until they reached relative confluence, and then passaged them at low split ratios (1:1, 1:1.5, or 1:2 dilutions) in complete organoid medium to promote continued growth. In one case, PANFR0489R, cells persisted as individuals and small organoids after initiation in complete organoid medium, but did not grow and expand cell numbers significantly. Approximately 15 weeks after initiation, we switched a portion of the surviving cells to organoid medium without A83-01 or mNoggin, and observed renewed growth of organoids under these media conditions but not of those that remained in complete organoid medium. Consequently, we expanded this sample in media without A83-01 or mNoggin, including performing early passage scRNA-seq. After several additional passages, once the organoids were robustly growing, we were able to transition back to complete organoid medium with no apparent change in growth rate, morphology, or transcriptional phenotype. All other serially sampled organoids were maintained and assessed by scRNA-seq in complete medium.

For scRNA-seq of organoid samples, we passaged organoids and allowed them to grow for 6 days before then dissociating, counting, and allocating 15,000 viable cells for Seq-Well. By standardizing the collection of organoid scRNA-seq samples at 6 days after passaging, we tried to minimize bias arising from cell cycle differences in samples at different degrees of confluence.

##### Testing organoid phenotypes under different matrix and media conditions

For adaptation of patient-derived organoids onto 2-dimensional (2D) culture surfaces as patient-derived cell lines, tissue culture plates were pre-coated with 100 μg/mL Matrigel dissolved in basal media for 2 hours at 37°C before washing with PBS. Established organoid models were dissociated and seeded onto these Matrigel-coated culture wells in complete organoid media. In parallel, a portion of these passage-matched organoid cells were re-seeded into Matrigel droplets as above. Cells were cultured in both matrix conditions in complete organoid media until they were confluent, approximately 2-3 weeks. Cells were collected and lysed using Trizol before snap freezing. RNA was isolated and purified as described below (“Bulk RNA-sequencing of organoids” section) using chloroform extraction, aqueous phase isolation, and processing using the Qiagen AllPrep DNA/RNA/miRNA Universal kit before being submitted for sequencing.

For scRNA-seq assessment of organoid phenotypes when cultured under different media conditions, established organoid models were passaged as above by dissociating and reseeding into Matrigel droplets. A portion of the cells were cultured with complete organoid media (“Complete media”), while a distinct portion of passage-matched cells were cultured in “Minimal” media, which consisted of advanced DMEM/F12 (Gibco), 10 mM HEPES (Gibco), 1x GlutaMAX (Gibco), 100 U/mL penicillin/streptomycin (Gibco), and 1x Primocin (Invivogen) (**Supplemental Table S6**). Cells were cultured for 6 days before being collected, dissociated, and aliquoted for scRNA-seq. Images were taken with an Olympus XM10 camera mounted to an Olympus CKX41 microscope 1 day after seeding and again after 11 days in culture to assess organoid growth in both conditions. The portion of cells cultured in minimal media were maintained in the same conditions for a longer duration and harvested again for scRNA-seq at 27 days and 59 days after the initial introduction of minimal media. To mirror the standard scRNA-seq workflow, the cells harvested at the 27- and 59-day timepoints were collected 6 days after passaging.

In addition to the minimal media experiment, organoid cells were also cultured in standard cell line media (“RP10”), which contains RPMI-1640 (Gibco) and 100 U/mL penicillin/streptomycin (Gibco) with 10% fetal bovine serum (Sigma), or in reduced organoid media “OWRNA”, which consists of advanced DMEM/F12 (Gibco), 10 mM HEPES (Gibco), 1x GlutaMAX (Gibco), 50 ng/mL mEGF (Peprotech), 100 ng/mL hFGF10 (Peprotech), 10 nM hGastrin I (Sigma), 1.25 mM N-acetylcysteine (Sigma), 10 mM Nicotinamide (Sigma), 1x B27 supplement (Gibco), 100 U/mL penicillin/streptomycin (Gibco), and 1x Primocin (Invivogen) (i.e. complete organoid medium with removal of WNT3A, RSPONDIN-1, NOGGIN, and A-8301; **Supplemental Table S6**). Furthermore, OWRNA reduced organoid medium served as the baseline control medium when assessing the effect of specific factors (IFNGγ and TGF-β1) from the TME on transcriptional phenotypes. Cells were cultured for 6 days before being collected, dissociated, and aliquoted for scRNA-seq in each of the following conditions: RP10, OWRNA, OWRNA with 50 ng/mL IFNGγ (Peprotech), and OWRNA with 5 ng/mL TGFB1 (Peprotech) (**Supplemental Table S6**). The cells cultured under the IFNGγ and TGF-β1 conditions were maintained in culture and harvested again for scRNA-seq 38 days after being introduced to these new media conditions. For these longer duration timepoints, cells were again passaged 6 days before collecting for scRNA-seq.

##### Testing transcriptional phenotype changes in an established cell line under organoid media conditions

For scRNA-seq assessment of transcriptional phenotypes of the established pancreatic cancer cell line CFPAC1 under different media conditions, CFPAC1 cells were cultured in parallel in either standard cell line medium RP10 or complete organoid medium. Cells were cultured for 6 days before being collected, dissociated, and aliquoted for scRNA-seq. Additionally, the CFPAC1 cells cultured under complete organoid medium were maintained in the same conditions and harvested again for scRNA-seq 33 days after the initial introduction of complete organoid medium. CFPAC1 cells grown in complete media for the later 33-day timepoint were collected 6 days after passaging, and media was refreshed 3 days after this final passage.

##### Single-cell RNA-seq (scRNA-seq) data library generation, sequencing, and alignment

ScRNA-seq processing followed the Seq-Well protocol, uniquely compatible with low-input samples (Gierahn et al., 2017; Hughes et al., 2020). Briefly, arrays were preloaded with RNA capture beads (ChemGenes) and stored in quenching buffer until used. Prior to cell loading, arrays were resuspended in 5 mL RPMI-1640 medium with 10% fetal bovine serum (both from Gibco, hereafter referred to as RP10). After dissociation, single-cell suspensions were manually counted and diluted to 15,000 cells per 200 μL of RP10 when cell numbers allowed. Excess RP10 was aspirated from the array and cells were loaded onto arrays. Excess cells were washed off with PBS (4×5 mL, Gibco), briefly left in RPMI (5 mL) and cell+bead pairs were sealed for 40 minutes at 37°C using a polycarbonate membrane (Fisher Scientific NC1421644). Arrays were rocked in lysis buffer for 20 minutes and RNA was hybridized onto the beads for 40 minutes. Beads were removed and reverse transcription was performed overnight using Maxima H Minus Reverse Transcriptase (Thermo Fisher EP0753). Prior to sequencing, the beads underwent an exonuclease treatment (NewEngland Biolabs M0293L) and second strand synthesis *en masse* followed by whole transcriptome amplification (WTA, Kapa Biosystems KK2602) in 1,500 bead reactions (50 μL). cDNA was isolated using Agencourt AMPure XP beads (Beckman Coulter, A63881) at 0.6X SPRI (solid-phase reversible immobilization) followed by a 1X SPRI and quantified using a Qubit dsDNA High Sensitivity assay kit (Thermo Fisher Q32854). Library preparation was performed using Nextera XT DNA tagmentation (Illumina FC-131-1096) and N700 and N500 indices specific to a given sample. Tagmented and amplified sequences were purified with a 0.6X SPRI. cDNA was loaded onto either an Illumina Nextseq (75 Cycle NextSeq500/550v2 kit) or Novaseq (100 Cycle NovaSeq6000S kit, Broad Institute Genomics Platform) at 2.4 pM. Regardless of platform, the paired end read structure was 21 bases (cell barcode and UMI) by 50 bases (transcriptomic information) with an 8 base pair (bp) custom read one primer. The demultiplex and alignment protocol was followed as previously described (Macosko et al., 2015). While Novaseq data were directly output as FASTQs, Nextseq BCL files were converted to FASTQs using bcl2fastq2. The resultant Nextseq and Novaseq FASTQs were demultiplexed by sample based on Nextera N700 and N500 indices. Reads were then aligned to the hg19 transcriptome using the cumulus/dropseq_tools pipeline on Terra maintained by the Broad Institute using standard settings.

##### Bulk RNA-sequencing of organoids

RNA was obtained for bulk RNA-sequencing from established organoids using one of two approaches. Dissociated organoids were resuspended into cold Matrigel, added as droplets to tissue culture plates (Greiner BioOne), and allowed to polymerize for 30 minutes before addition of media. Organoids were grown for 14-21 days (until confluent) under these conditions with regular media changes. At the time of harvest, cells were washed with cold phosphate buffered saline (PBS) at 4°C, then lysed with Trizol (Invitrogen) before snap-freezing. To isolate RNA, we performed chloroform extraction with isolation of the aqueous phase before processing RNA as per protocols outlined in the Qiagen AllPrep DNA/RNA/miRNA Universal kit.

In the second approach, dissociated organoids were resuspended in a solution of 10% Matrigel in complete organoid media (volume/volume) and cultured in ultra-low-attachment culture flasks (Corning). Organoids were grown for 14-21 days (until confluent) before pelleting, washing with cold PBS at 4°C until most Matrigel was dissipated, and then snap frozen. For RNA isolation, cell pellets were homogenized using buffer RLT Plus (Qiagen) and a Precellys homogenizer. Samples were then processed for both DNA extraction and RNA isolation as per the Qiagen AllPrep DNA/RNA/miRNA Universal kit. Purified RNA was then submitted for sequencing by the Broad Institute Genomics Platform.

In brief, total RNA was quantified using the Quant-iT RiboGreen RNA Assay Kit (Thermo Fisher R11490) and normalized to 5 ng/μL. Following plating, 2 μL of a 1:1000 dilution of ERCC RNA controls (Thermo Fisher 4456740) were spiked into each sample. An aliquot of 200 ng for each sample was transferred into library preparation which uses an automated variant of the Illumina TruSeq Stranded mRNA Sample Preparation Kit. This method preserves strand orientation of the RNA transcript, and uses oligo dT beads to select mRNA from the total RNA sample followed by heat fragmentation and cDNA synthesis from the RNA template. The resultant 400 bp cDNA then goes through dual-indexed library preparation: ‘A’ base addition, adapter ligation using P7 adapters, and PCR enrichment using P5 adapters. After enrichment, the libraries were quantified using Quant-iT PicoGreen (1:200 dilution; Thermo Fisher P11496). After normalizing samples to 5 ng/μL, the set was pooled and quantified using the KAPA Library Quantification Kit for Illumina Sequencing Platforms. The entire process was performed in 96-well format and all pipetting was done by either Agilent Bravo or Hamilton Starlet.

Pooled libraries were normalized to 2 nM and denatured using 0.1 N NaOH prior to sequencing. Flowcell cluster amplification and sequencing were performed according to the manufacturer’s protocols using the NovaSeq 6000. Each run was a 101 bp paired-end with an eight-base index barcode read. Data were analyzed using the Broad Picard Pipeline which includes de-multiplexing and data aggregation (https://broadinstitute.github.io/picard/). FASTQ files were then processed as described below (see Bulk RNA-sequencing analysis).

##### Multiplex immunofluorescence imaging

A multi-marker panel was developed to characterize tumor cell subtype in formalin-fixed paraffin-embedded (FFPE) 4μm tissue sections using multiplex immunofluorescence. The panel comprises markers associated with either a basal (Keratin-17: Thermo Fisher MA513539 and s100a2: Abcam 109494) or classical (cldn18.2: Abcam 241330, GATA6: CST 5851 and TFF1: Abcam 92377) subtype. Additionally, DAPI (Akoya Biosciences FP1490) was included for identification of nuclei and pan-cytokeratin (AE1/AE3: DAKO M3515; C11: CST 4545) for identification of epithelial cells. Secondary Opal Polymer HRP mouse and rabbit (ARH1001EA), Tyramide signal amplification and Opal fluorophores (Akoya Biosciences) were used to detect primary antibodies (Keratin-17, Opal 520; s100a2, Opal 650; GATA6, Opal 540; cldn18.2, Opal 570; TFF1, Opal 690; panCK, Opal 620). Prior to use in multiplex staining, primary antibodies were first optimized via immunohistochemistry on control tissue to confirm contextual specificity. Monoplex immunofluorescence and iterative multiplex fluorescent staining were then used to optimize staining order, antibody-fluorophore assignments and fluorophore concentrations. Multiplex staining was performed using a Leica BOND RX Research Stainer (Leica Biosystems, Buffalo, IL) with sequential cycles of antigen retrieval, protein blocking, primary antibody incubation, secondary antibody incubation, and fluorescent labeling. Overview images of stained slides were acquired at 10X magnification using a Vectra 3.0 Automated Quantitative Imaging System (Perkin Elmer, Waltham, MA) and regions of interest (ROIs) were selected for multispectral image acquisition at 20X. After unmixing using a spectral library of single-color references, each image was inspected to ensure uniform staining quality and adequate tumor representation.

#### Data analysis

##### Mutation and CNV identification from bulk DNA-sequencing

For targeted DNA-sequencing of clinical samples, next-generation sequencing using a custom-designed hybrid capture library preparation was performed on an Illumina HiSeq 2500 with 2×100 paired-end reads, as previously described (Garcia et al., 2017; Sholl et al., 2016). Sequence reads were aligned to reference sequence b37 edition from the Human Genome Reference Consortium using bwa, and further processed using Picard (version 1.90, http://broadinstitute.github.io/picard/) to remove duplicates and Genome Analysis Toolkit (GATK, version 1.6-5-g557da77) to perform localized realignment around indel sites. Single nucleotide variants were called using MuTect v1.1.45, insertions and deletions were called using GATK Indelocator. Copy number variants and structural variants were called using the internally-developed algorithms RobustCNV and BreaKmer followed by manual review (Abo et al., 2015). RobustCNV calculates copy ratios by performing a robust linear regression against a panel of normal samples. The data were segmented using circular binary segmentation, and event identification was performed based on the observed variance of the data points (Bi et al., 2017).

We computed the cytoband-level copy number calls and weighted (by length) average segment means across the covered regions of each cytoband using ASCETS (Spurr et al., 2020). Briefly, cytobands were considered amplified/deleted if more than 70% of the covered regions had a log2 copy ratio of greater than 0.2/less than −0.2, and were considered neutral if more than 70% of the covered regions had a log2 copy ratio between −0.2 and 0.2.

##### Single-cell data quality pre-processing and initial cell type discovery

All single-cell data analysis was performed using the R language for Statistical Computing (v3.5.1). Each biopsy sample’s digital gene expression (DGE) matrix (cells x genes) was trimmed to exclude low quality cells (<400 genes detected; <1,000 UMIs; >50% mitochondrial reads) before being merged together (preserving all unique genes) to create the larger biopsy dataset. The merged dataset was further trimmed to remove cells with >8,000 genes which represent outliers and likely doublet cells. We also removed genes that were not detected in at least 50 cells. The same metrics were applied to the organoid single-cell cohort (see below). On a per cell basis, UMI count data was divided by total transcripts captured and multiplied by a scaling factor of 10,000. These normalized values were then natural log transformed for downstream analysis (i.e. log-normalized cell x gene matrix). Initial exploration of the data was performed using the R package Seurat (v2.3.4) and followed two steps: 1) SNN-guided quality assessment and 2) cell type composition determination. In step 1, we intentionally left cells in the DGE matrix of dubious quality (e.g. % mitochondrial reads >25% but <50%), performed principal component analysis (PCA) over the variable genes (n = 1,070 genes), and input the first 50 PCs (determined by Jackstraw analysis implemented through Seurat) to build an SNN graph and cluster the cells (res = 1; k.param = 40). The inclusion of poor-quality cells essentially acts as a variance “sink” for other poor-quality cells and they cluster together based on their shared patterns in quality-associated gene expression. This method helped to nominate additional low quality (e.g. defined exclusively by mitochondrial genes) or likely doublet cells (e.g. clusters defined by co-expression of divergent lineage markers) which were removed from the dataset (n=1,678 cells). This led to an overall high-quality dataset of single-cells with a low overall faction of mitochondrial reads (median = 0.09) for downstream analysis (**Supplemental Figure S1B**)

Using the trimmed dataset, we proceeded to step 2 using a very similar workflow as above but with slightly altered input conditions for defining clusters. Here we used PCs 1-45 and their associated statistically significant genes for building the SNN graph and determining cluster membership (resolution = 1.2; k.param = 40). This identified the 36 clusters shown (visualized using *t*-SNE; perplexity, 40; iterations, 2,500) in **Supplemental Figure S1C**. The expression of known markers was used to collapse clusters containing shared lineage information. For example, clusters 1, 2, and 4 all express high levels of macrophage markers—*CD14*, *FCGR3A (CD16)*, *CD68*—and were accordingly collapsed for this first pass analysis (**Supplemental Figure S1C,G**). To aid our cell type identification, we performed a ROC test implemented in Seurat to confirm the specificity (power > 0.6) of the top marker genes used to discern the cell types. Combined with inferred CNV information (see below), this analysis confirmed the presence of 11 broad non-malignant cell types in our biopsy dataset (**Supplemental Table S2**). Variation in the SNN graph parameters above did not strongly affect cell type identification.

##### Single-cell CNV identification

To confirm the identity of the putative malignant clusters identified in **Supplemental Figure S1D**, we estimated single-cell CNVs as previously described by computing the average expression in a sliding window of 100 genes within each chromosome after sorting the detected genes by their chromosomal coordinates (Patel et al., 2014; Tirosh et al., 2016b). We used all T/NK, Fib, Hep, and Endo cells identified above as reference normal populations for this analysis. Complete information on the inferCNV workflow used for this analysis can be found here https://github.com/broadinstitute/inferCNV/wiki. To compare with bulk targeted DNA-sequencing, we collapsed individual probes to cytoband-level information (weighted average of log2 ratios across each cytoband, see above) within each sample. ScRNA-seq-inferred CNVs showed high concordance across samples with the bulk measurements and suggests that, at least by this metric, we are likely sampling the same dominant clones within sequential but distinct cores from each needle biopsy procedure (**Supplemental Figure S1E**). For plotting CNV profiles in putative malignant versus normal cells (**Supplemental Figure S1F**), we computed the average CNV signal for the top 5% of altered cells in each biopsy and correlated all cells in that biopsy to the averaged profile as has been previously described (Tirosh et al., 2016a). Relation of this correlation coefficient to the CNV score (mean square deviation from diploidy) in the single cells from each biopsy shows consistent separation of malignant from non-malignant cells, and, combined with membership in patient-specific SNN clusters, substantiates the identification of malignant cells in our dataset.

##### Subclonal analysis with single-cell inferred CNVs

The inferCNV workflow can be used to call subclonal genetic variation with high sensitivity and is comprehensively outlined here https://github.com/broadinstitute/inferCNV/wiki (Fan et al., 2018; Patel et al., 2014; Tirosh et al., 2016b). Briefly, we used a six-state Hidden Markov Model (i6-HMM) to predict relative copy number status (complete loss to >3x gain) across putative altered regions in each cell. A Bayesian latent mixture model then evaluated the posterior probability that a given copy number alteration is a true positive. We set a relatively stringent cutoff for this step (BayesMaxPNormal = 0.2) to only include high probability alterations for downstream clustering. The results of this filtered i6-HMM output were then used to cluster the single cells using Ward’s method. We used inferCNV’s “random trees” method to test for statistical significance (*P* < 0.05, 100 random permutations for each split) at each tree bifurcation and only retained subclusters that had statistical evidence underlying the presumed heterogeneity. To track subclonal heterogeneity between biopsy and matched organoid cells in **Figure 3G** and **Supplemental Figure S5E-I**, the above workflow was implemented within each biopsy and the relevant matched organoid samples, essentially treating all cells as the same “tumor” and allowing the CNVs to determine cell sorting agnostic to sample-of-origin. The results of the HMM output can be used to infer gene-level information based on which genes are in the affected window. This (like the rest of the HMM workflow) is computed over groups of cells (e.g. samples or sub-clones) and used to map *KRAS* and other alterations to samples (**Figure 3A-F**) or sub-clones (**Figure 3G, Supplemental Figure S5E-I**).

##### Subclustering of malignant and non-malignant cells

Detailed phenotyping required splitting the dataset into malignant and non-malignant fractions. After subsetting to only the malignant cells, we re-scaled the data and ran PCA including the first 35 PCs for SNN clustering and *t-*SNE visualization. This PCA was used to determine the PanNET identity for PANFR0580 (**Supplemental Figure S2A**). After removing PANFR0580, we repeated the steps above and used this new PCA for the remainder of PDAC malignant cell analysis. Subsequent phenotyping for malignant cells is discussed below (**Generation of expression signatures/scores**). A similar approach was used for calling the non-malignant subsets in **Figure 5A**. To determine the specific phenotypes within T/NK, macrophage, and mesenchymal populations, we separately subclustered these groups using PCs 1-20 and a resolution of 0.6 in each case. Of note, subclustering within the macrophages revealed a distinct cluster of cells co-expressing markers of both T/NK cells and macrophages (n=491 cells). We discarded these cells as likely doublets, as have others, and re-ran the macrophage PCA and clustering (Zhang et al., 2020; Zilionis et al., 2019). These cells are included in the full dataset in case they are of interest to others. Each unbiased analysis helped to define the non-malignant phenotypes summarized in **Figures 5 & 6** and **Supplemental Figure S6**.

##### Generation of expression signatures/scores

All expression scores were computed as previously described by taking a given input set of genes and comparing their average relative expression to that of a control set (n=100 genes) randomly sampled to mirror the expression distribution of the genes used for the input (Tirosh et al., 2016b). While all scores were computed in the same way, choosing the genes for input varied. We have outlined the relevant approaches below. Where correlations (Pearson’s *r)* are performed over genes, we used the log-transformed UMI count data for each case. Unless otherwise noted, we selected the top 30 statistically significant genes for each signature (>3 s.d. above the mean for shuffled data) for visualization and scoring.

##### Cell cycle

We utilized previously established signatures for G1/S (n=43 genes) and G2/M (n=55 genes) to place each cell along this dynamic process (Tirosh et al., 2016a). After inspecting the distribution of scores in the complete dataset, we considered any cell >1.5 s.d. above the mean for either the G1/S or the G2/M scores to be cycling (van Galen et al., 2019).

##### Basal and classical programs

We started by scoring each malignant single cell for the basal-like and classical genes identified by Moffitt et al., 2015 as these were well described by unbiased analysis in our data (PCA, **Supplemental Figure S2B**). To determine programs associated with basal and classical phenotypes, we correlated the aforementioned basal and classical scores to the entire gene expression matrix containing malignant cells and selected the 1,909 genes significantly associated with either subtype (*r* > 0.1; >3 s.d. above the mean for shuffled data, full data in **Supplemental Table S3**). Biological pathway correlates for basal and classical mirrored previous work, and are summarized in **Supplemental Figure S3D,E**. For visualization, we use the “scCorr” basal and classical genes (top 30 correlated genes for each). We used these basal and classical scores to order the cells by their polarization or “score difference”, simply the difference of the two scores, and revealed a significant fraction of cells co-expressing intermediate levels of both phenotypes (**Supplemental Figure S3A,B**).

##### Intermediate transitional program

Intermediate cells showed associations with features across several additional PCs, but lacked a single dominant axis. To define a consensus set of genes that are preferentially expressed by cells in this intermediate state, we computed the Euclidean distance to the line representing equal basal and classical co-expression for each cell. To limit the influence of cell quality on this analysis and to specifically identify genes related to co-expression, we used cells from each group (basal, intermediate, and classical) with fractionally low mitochondrial genes (<0.2) and non-zero basal or classical expression (basal or classical score > 0) and correlated their Euclidean distance (**Supplemental Figure S3C**) to the entire gene expression matrix of malignant cells. Next, for each gene positively associated with this intermediate state (Pearson’s *r* >0), we subtracted the second highest correlation coefficient for each subtype-associated gene (basal and classical), and then re-ranked the matrix by this corrected value. This enriched for genes more specific to the intermediate state by excluding those that were also associated with basal or classical programs. We then selected the 115 genes with a corrected correlation value >0.1 (*P< 0.00001*, shuffled data) as our intermediate transitional (IT) signature (**Supplemental Figure S3D**, **Supplemental Table S3**). Single cells were classified based on Euclidian distance where <0.2 are defined as intermediate transitional and the remainder (Euclidian distance >0.2) by their maximal of either basal or classical scores. We binned each organoid cell (e.g. **Figure 4B,C**) by its maximal expression for one of the 3 *in vivo* scores (basal, classical, or IT). Here a cell must be within 1 s.d. of the mean expression for a given subtype *in vivo*, else it was considered “organoid-specific” as this program was superimposed on all organoid cells, regardless of their subtype identity (**Figure 4B**). We used these classifications to summarize overall tumor composition and visualize the groups. Tumor heterogeneity measures were not significantly affected by changing these cutoffs.

##### Non-Malignant programs

TAM signatures were determined similar to above and previous work (van Galen et al., 2019; Zhang et al., 2020; Zilionis et al., 2019). Using PCA as an anchor (**Supplemental Figure S6C**), we correlated expression within the TAM compartment to either *FCN1*, *SPP1*, or *C1QC* (top loaded genes on each relevant PC) and merged the resultant correlation coefficients for every detected gene to the 3 subtypes into one matrix (i.e. a 16,920 x 3 matrix). For each TAM type (i.e. vector of correlation coeffects to each marker), we first ranked the matrix by decreasing correlation coefficient, selected only the most significantly associated genes to that type (*r* > 0.1; >3 s.d. above the mean for shuffled data), subtracted the second highest correlation coefficient for each subtype-associated gene, and then re-ranked the matrix by this corrected value. We repeated this procedure for each TAM subtype independently. This ensures that the genes selected are specific to a given TAM subset and do not describe general TAM features. The top 30 genes for each were used for scoring and visualization (**Supplemental Table S2**; **Supplemental Figure S6D**).

CAF phenotypes were determined using a similar workflow. To examine fibroblast heterogeneity, we removed a subset of adrenal endocrine cells (cluster 4, 40 cells; **Figure 5C**) and then performed PCA of mesenchymal cells. PC1 was driven by spillover genes (likely contributed from ambient RNA) and lacked any coherent biological program and was not considered further. PCs 2 and 3 by contrast where consistent with variable mesenchymal (PC2) and inflammatory (PC3) CAF phenotypes. All these cells scored highly for previous myCAF gene expression programs so this phenotype did not fully explain the heterogeneity in mesenchymal cells, but did suggest their identity as CAFs. Again, using correlation, we determined the genes driving low PC2 scores (Dermal-like), and high PC2 scores (Pericyte-like), as well as those associated with the high PC3 scores (Inflammatory). As before, we used the top 30 genes for each subset scoring and visualization. These same genes (Dermal-like and Pericyte-like) were used to examine bulk RNA-seq profiles and their difference in each sample quantifies which phenotype is favored in the bulk averages (**Figure 5F**).

##### TME associations

We determined the transcriptional-subtype-dependent composition of the TME (**Figure 6A-C**) following two steps. First, we computed the Simpson’s Index (measure of ecological diversity) using the count of each non-malignant cell type captured from each sample as input (**Figure 6A,B**) and correlated each biopsy’s diversity score to its basal vs. classical commitment score. Importantly, the number of non-malignant cells captured from each biopsy was not associated with basal vs. classical commitment score (r = 0.09). Next, to understand which cell types drive these differences, we computed the fractional representation for every non-malignant cell type in each core needle biopsy and determined their pairwise correlation distance (Pearson’s *r*) followed by hierarchical clustering using Ward’s method (dendrogram in **Figure 6B**). For both of these analyses we only used samples with >200 non-malignant cells captured (**Supplemental Figure S6L**).

##### Matched organoid clustering and cell-typing

After applying similar quality metrics as above, we performed PCA, SNN clustering, and *t*-SNE embedding for 31,867 cells including organoid cells and all malignant cells from primary PDAC biopsies (PCs 1-50; resolution=1.2; k.param=45; perplexity=45; max_iter=2,500), and identified 39 total clusters. Organoids clustered separately from their matched biopsies, suggesting expression and/or CNV related drift in culture. Only two SNN clusters—clusters 4 and 32—were admixed by sample. We determined the specific gene expression programs in these two clusters via differential expression testing by Wilcoxon rank sum test (*P* < 0.05, Bonferroni correction; log(fold change) > 0.5). These comparisons were done in a “1 versus rest” fashion, testing for genes defining each cluster (4 or 32) compared to the entire dataset. Their expression profiles were consistent with fibroblasts (cluster 32) and epithelial cells (cluster 4; **Supplemental Figure S5B,C**).

##### Correlation distances for genotype and phenotype

To generate correlation distances for genotype and phenotype, each single cell in a biopsy-organoid pair was represented by two vectors of information: (i) a phenotype vector containing expression values for basal and classical genes (scCorr basal and classical genes, n = 60 genes) and (ii) a genotype vector containing the average CNV score for each cytoband. The phenotype and genotype distances between every single cell within a biopsy/early organoid pair was computed from these vectors using a correlation-based (Pearson’s *r*) distance metric of the form *d* = (1-*r*)/2. This resulted in two distance matrices of n x n dimension where n is the total number of cells from each biopsy/early organoid sample pair. Values in **Figure 4A** are computed by averaging the values for *d* between only early organoid and matched biopsy cells.

##### Matched biopsy vs. organoid malignant cell comparison

For CNV-confirmed malignant cells from each biopsy and its matched organoid (earliest passage), we used differential expression (Wilcoxon rank sum test; *P* < 0.05, Bonferroni correction; log(fold change) > 0.3) to understand the features lost from malignant cells in the *in vivo* setting and gained when transitioning into growth in organoid culture. We required any gene to be significantly differentially expressed in at least 3 model-biopsy comparisons to summarize the consistent changes. We repeated this same workflow for both organoid- and biopsy-specific genes (**Supplemental Table S5**) outlined in **Figure 4B** and **Figure 4D-F**, respectively.

##### Biopsy paracrine and autocrine subtype-specific factor analysis

Factors present in the TME but absent from organoid culture could originate from at least two sources, the tumor cells themselves (autocrine) or non-tumor cells in the local microenvironment (paracrine). We examined any gene with gene ontology annotations related to “cytokines”, “chemokines”, or “growth factors” and took the union of these lists, yielding 321 genes, 218 of which were detected in our dataset. For “autocrine” factors we performed differential expression between malignant cells binned as basal and classical, and then IT vs rest. A gene was considered differentially expressed if it passed a *P* < 0.05 with Bonferroni correction and a log(fold change) > 0.2 in one of these comparisons. Genes were then assigned to subtypes based on the log fold change direction (**Figure 7D**, **Supplemental Table S7**). Paracrine factors were determined in a similar manner with slight modifications. We grouped non-tumor cells into basal, classical or IT based on the average expression and clustering for malignant programs from their respective tumor samples (**Supplemental Figure S3G,H**). We then assessed for differential expression between all cells from a given group and the rest using the same cutoffs as above and sorted factors into subtypes based on their log fold change directionality (**Figure 7F**, **Supplemental Table S7**). We then visualized which cell type contributed the highest average expression for each factor in the cell types from the respective TMEs (**Figure 7G**).

##### Bulk RNA-sequencing analysis

FASTQs for bulk RNA expression profiles were downloaded from the relevant repository (TCGA, https://toil.xenahubs.net; PDAC Cell lines, https://portals.broadinstitute.org/ccle), available in-house (Panc-Seq, metastatic PDAC), or generated for this study (organoid cohort) (Aguirre et al., 2018; Cancer Genome Atlas Research et al., 2013; Cancer Genome Atlas Research Network, 2017; Ghandi et al., 2019). All were processed using the same pipeline. Briefly, each sample’s sequences were marked for duplicates and then mapped to hg38 using STAR. After running QC checks using RNAseqQC, gene-level count matrices were generated using RSEM. Instructions to run the pipeline are given in the Broad CCLE github repository https://github.com/broadinstitute/ccle_processing. Length-normalized values (TPM) were then transformed according to log2(TPM+1) for downstream analysis. The entire dataset was scaled and centered to allow relative comparisons across sample types (e.g. tumors, organoids, and cell lines). Signature scores were computed as above (e.g. basal and classical; see **Generation of expression signatures/scores** above) (Puram et al., 2017).

##### Tumor phenotyping from mIF data

Supervised machine learning algorithms were applied for tissue and cell segmentation (inForm 2.4.1, Akoya Biosciences). Single-cell-level imaging data were exported and further processed and analyzed using R (v3.6.2). To assign phenotypes to individual tumor epithelial cells, mean expression intensity in the relevant subcellular compartment was first used to classify cells as positive or negative for each of the 5 markers. Combinatorial expression patterns for the five markers were then used to phenotypically classify cells as basal, classical, co-expressing / IT or marker negative (3 combinations of 2 basal markers, 7 combinations of 3 classical markers, 1 pan-marker negative, 21 combinations of co-expression of basal and classical markers, **Supplemental Figure S4A, Supplemental Table S4**). Tumor subtype composition was assessed by calculating the fraction of total tumor cells positive for each cell phenotype (**Supplemental Figure S4B**, excluding pan-marker negative cells).

## SUPPLEMENTAL INFORMATION

**Supplemental Table S1, related to Figure 1.**
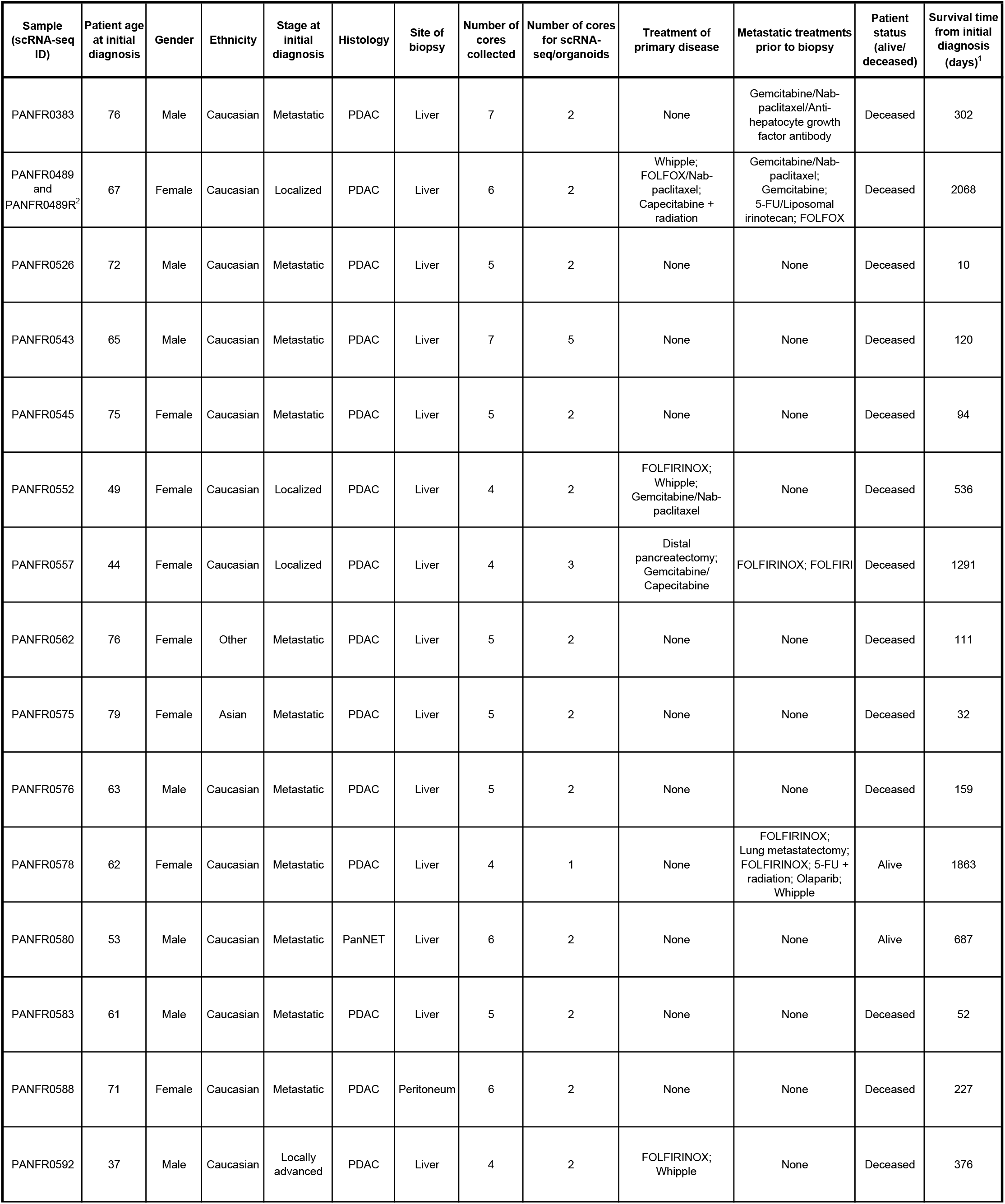
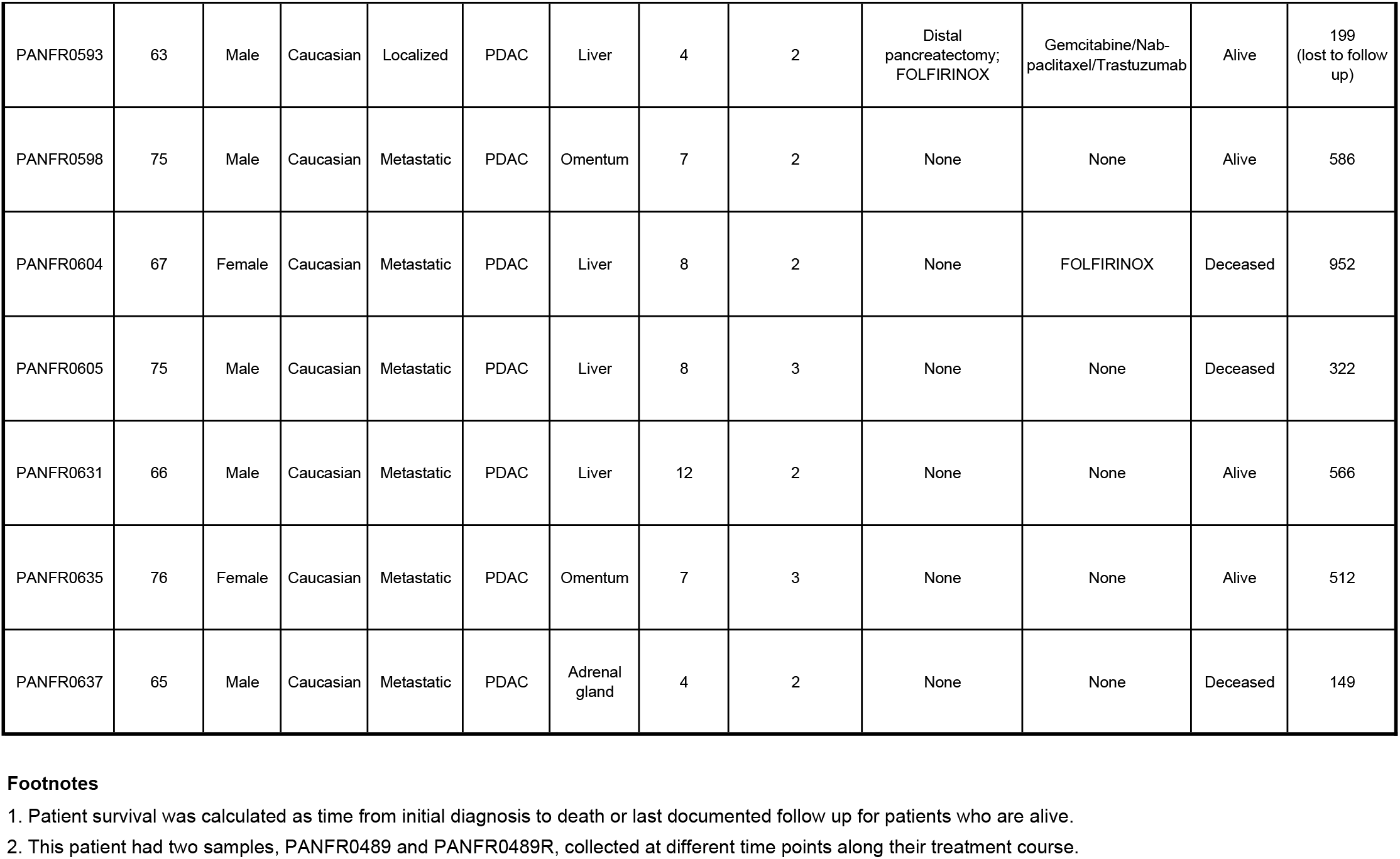
**Cohort patient characteristics** This table details the demographic and clinical characteristics of patients whose biopsy samples were used in this study.

**Supplemental Table S2. Normal cell type markers.**

*Related to Figures 1, 5 & 6*

**Supplemental Table S3. Malignant phenotype single-cell gene correlates.**

*Related to Figure 2*

**Supplemental Table S4. mIF marker combinations and cell counts.**

*Related to Figure 2*

**Supplemental Table S5. Organoid- and in vivo malignant-specific gene expression features.**

*Related to Figure 4*

**Supplemental Table S6. Organoid and cell line models and media formulations for perturbation experiments.**

*Related to Figure 7*

**Supplemental Table S7. Subtype-specific autocrine and paracrine secreted factors.**

*Related to Figure 7*

**Supplemental Figure S1.**
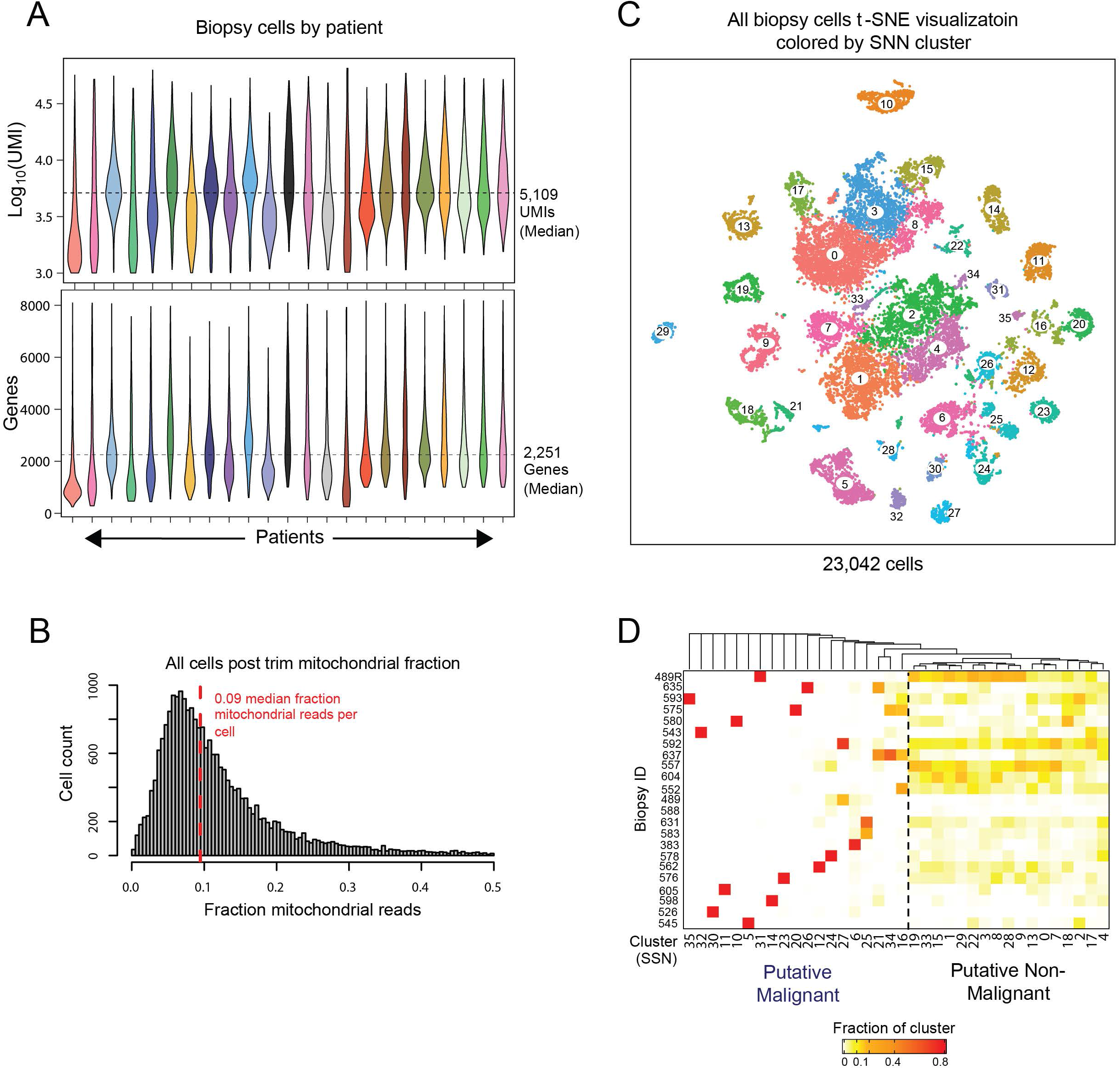
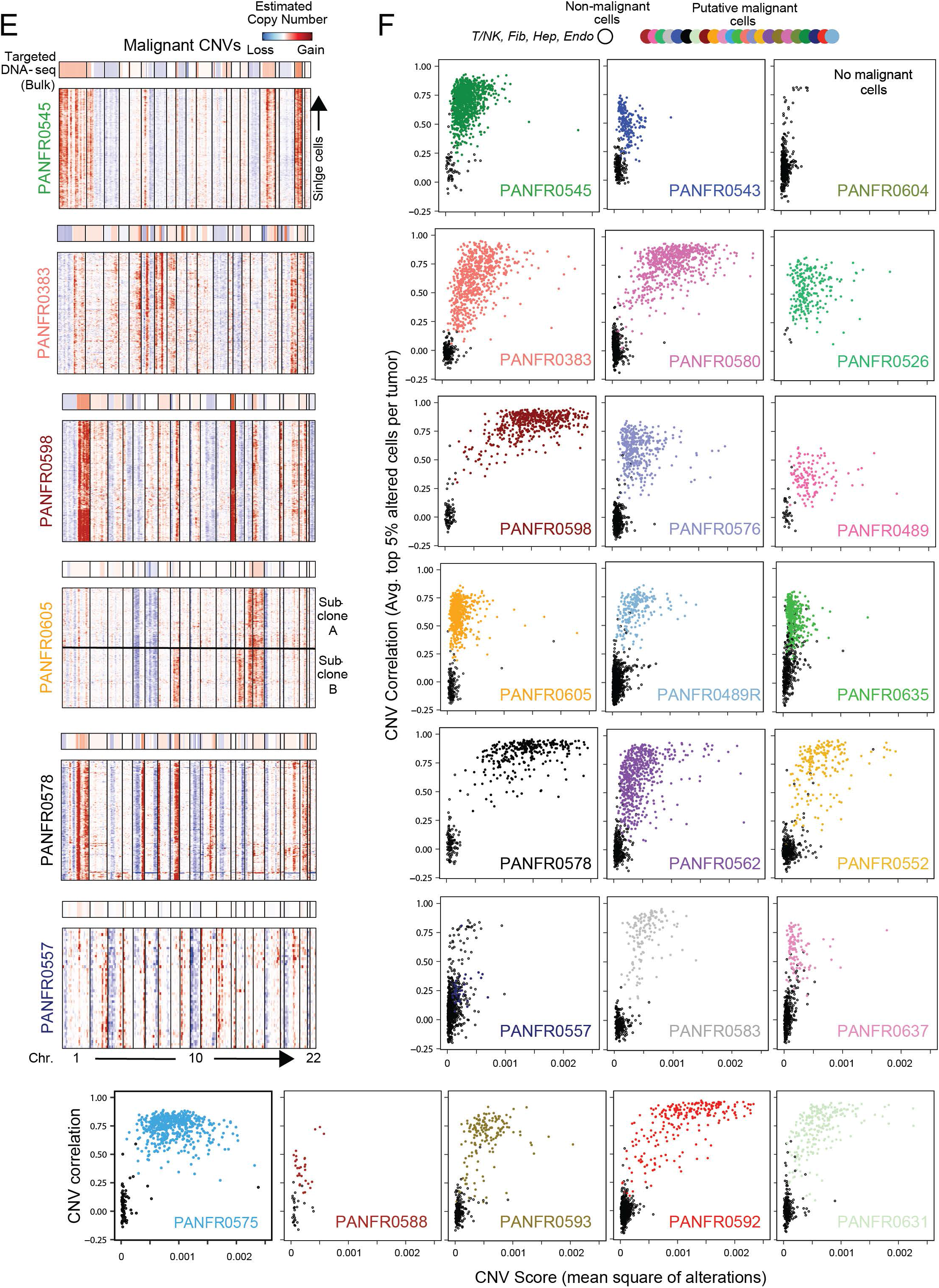
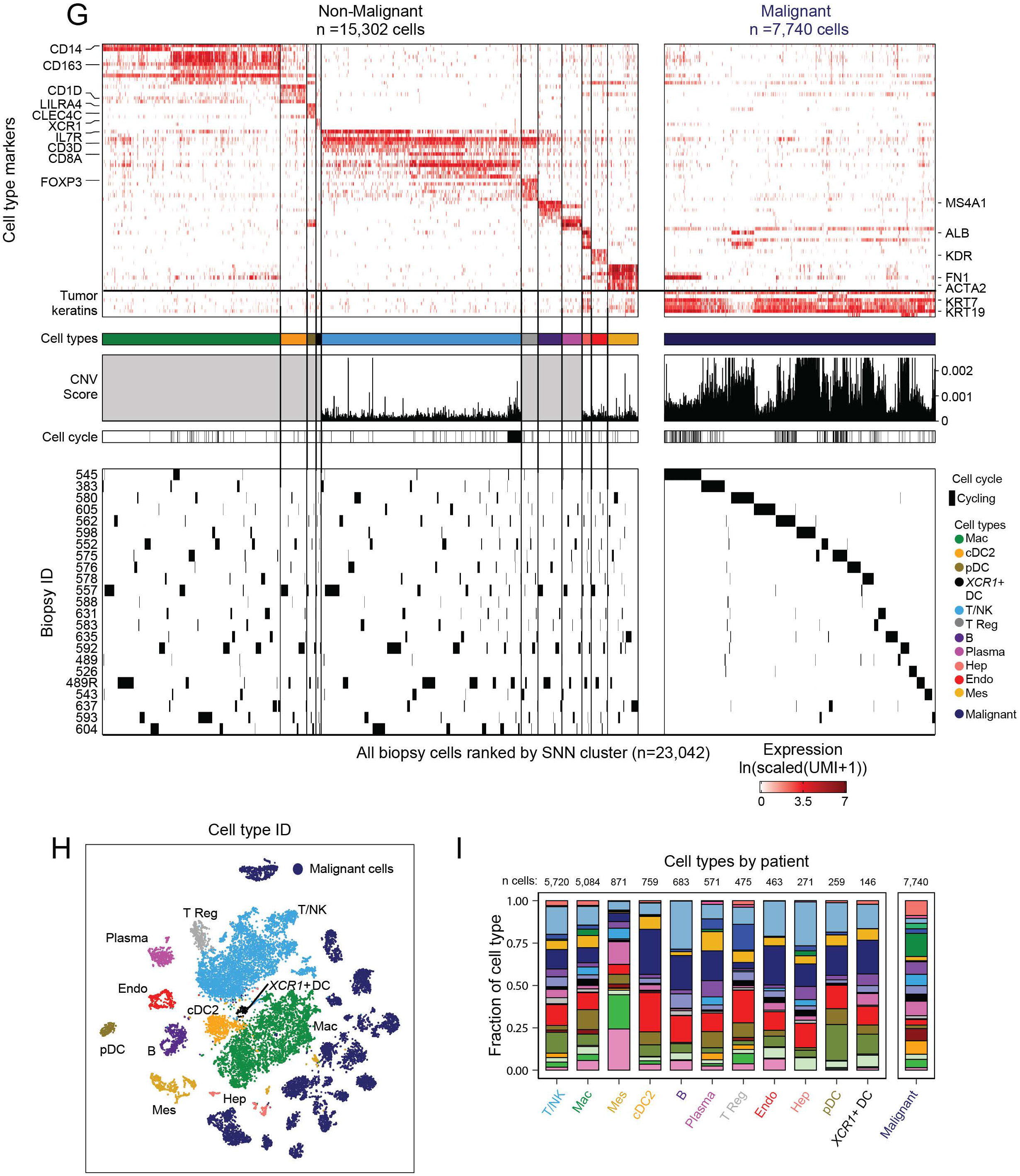
Quality metrics, unsupervised cell type identification, and malignant cell confirmation across the biopsy cohort. *Related to Figure 1* **A**, Distribution of unique molecules and genes captured in quality cells per biopsy, median values are indicated for each metric (dotted line) and violin plots are colored by patient (top, Log10(UMIs); bottom, number of genes). **B**, Distribution of fraction mitochondrial reads across the entire trimmed biopsy dataset (n = 23,042 cells). Red dotted line denotes the median. **C**, *t-*SNE visualization of the entire single-cell biopsy dataset colored by the SNN clusters identified (inset numbers). **D**, Distribution of single cells captured per biopsy across the identified SNN clusters. In general, a patient’s malignant cells are expected to form unique clusters driven by CNVs. Owing to this feature, the data are split into putative malignant and non-malignant groups of clusters. **E**, Heatmaps represent select scRNA-seq-derived copy number profiles where expression across the transcriptome is organized by chromosome (columns) for each single putative malignant cell (rows) from a given biopsy. Top bar indicates reference bulk targeted DNA-seq for the same patient and shows strong concordance with the single-cell derived profiles. **F**, CNV correlation (averaged top 5% of altered cells per biopsy) versus CNV score (mean square of modified expression) for each single putative malignant (colored points) and reference normal cell (empty black circles) within a given biopsy. Only a single sample, PANFR0604, did not contain any malignant cells. **G**, Overview of cell-typing for all cells in the biopsy dataset. Cells are ordered by SNN cluster and separated by cell types. Top heatmap represent expression levels for a subset of select markers (n=73 genes) used to identify cell types. Color bar indicates cell types and binarized cell cycle phenotypes are labeled (black, cycling; white, not). CNV scores (mean square of alterations per cell) used to parse malignant from non-malignant are shown using T/NK, endothelial, fibroblasts, and hepatocytes as reference; grey boxes denote normal cell types where we did not compute reference CNV scores. Bottom panel shows biopsy of origin for each cell. The data are split by non-malignant (n = 15,302) and malignant (7,740) identity. **H**, *t-*SNE visualization as in **S1C** but colored by cell types identified, abbreviations as in **Figure 1D**. **I**, Fraction of each cell type contributed by each biopsy sample (color fill, patient ID; as in **Figure 1B**), cell type totals are noted at the top of each bar.

**Supplemental Figure S2.**
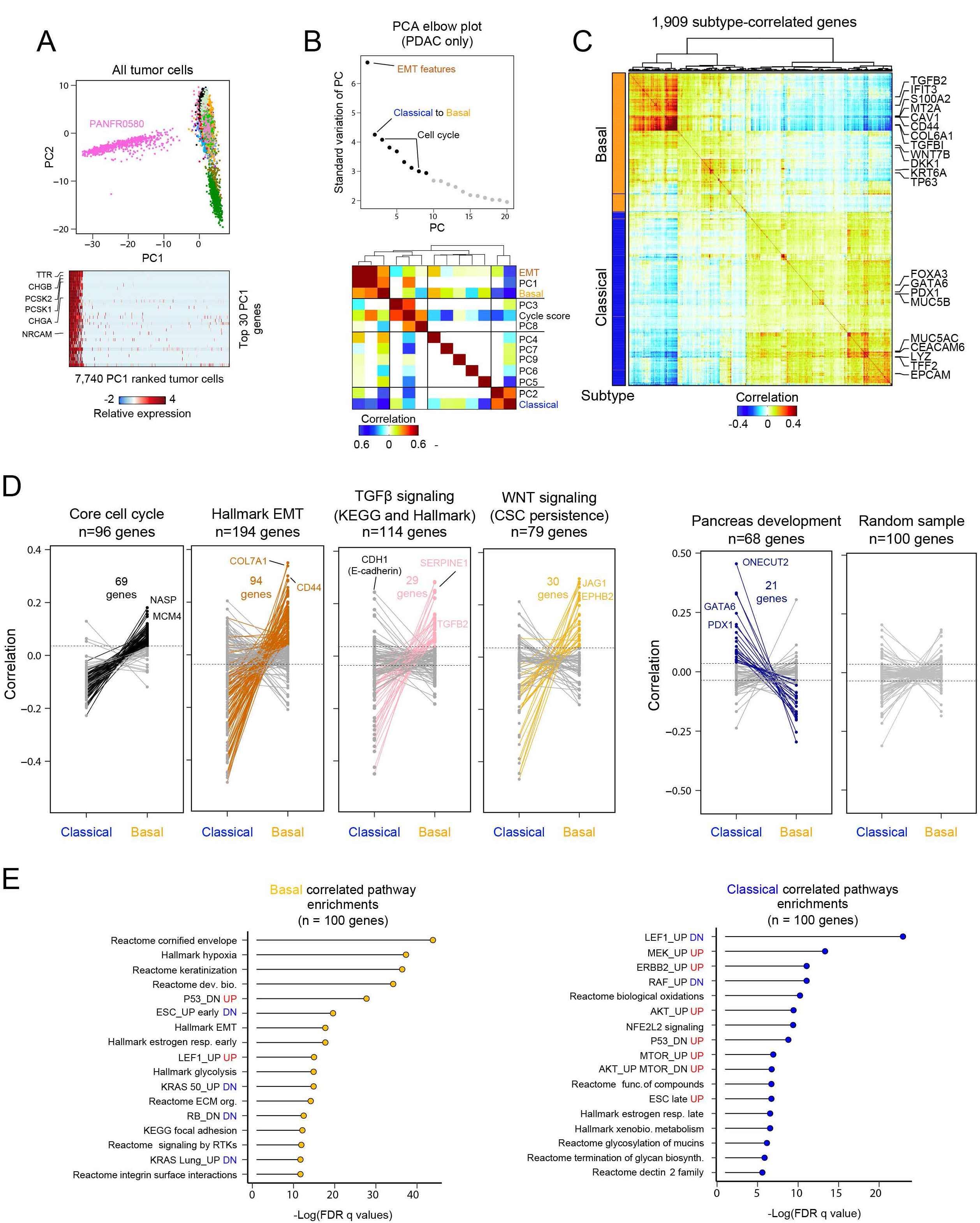
Identifying and contextualizing basal and classical associated biology. *Related to Figures 1 & 2* **A**, Principal component analysis (PCA) and scatter plot for PC1 and PC2 across all malignant cells (n=7,740) separates PANFR0580’s malignant cells (n=662) from the rest of the samples. Cells are colored by patient ID (as in **Figure 1B**). Heatmap for genes with the strongest negative loading on PC1 (n=30) denote a neuroendocrine identity (*TTR, CHGB*). This tumor was later classified by histology as a pancreatic neuroendocrine tumor (PanNET). **B**, Principal component (PC) elbow plot showing the standard deviation for the first 20 components calculated over the verified PDAC malignant cell variable genes (**Methods**). Line is drawn at the putative “elbow” (black versus grey points) as inclusion of additional PCs described overlapping information or quality metrics. Cross-correlational analysis for each single-cell’s embeddings across first 9 PCs (black points) and scores for literature curated gene sets describing EMT, classical and basal, and cell cycle phenotypes. PC1 positively correlates with EMT, basal, and to a degree, cell cycling. Cells with positive embeddings on PC2 are correlated with classical phenotypes and anti-correlated with basal and EMT phenotypes, suggesting these phenotypes are anti-correlated across a continuum of expression. PC3 and PC8 describe cells with high cell cycle scores. The other PCs do not associate significantly with these phenotypes. **C**, Pairwise correlation of genes significantly associated with basal (PC1/negative PC2) or classical (PC2) expression states. Left bar indicates the subtype association of each gene (orange, basal; blue, classical). **D**, Tied dot plots depicting the correlation coefficient for each gene (points) to either basal or classical phenotypes from select literature-derived gene sets, indicated at the top of each plot, which summarize aspects of subtype associated biology. Dotted lines represent significance threshold (3 SD above the mean of shuffled data), points and lines are colored if that gene passes the threshold and select genes are indicated. **E**, GSEA pathway enrichments for top 100 genes correlated to either basal or classical expression scores.

**Supplemental Figure S3.**
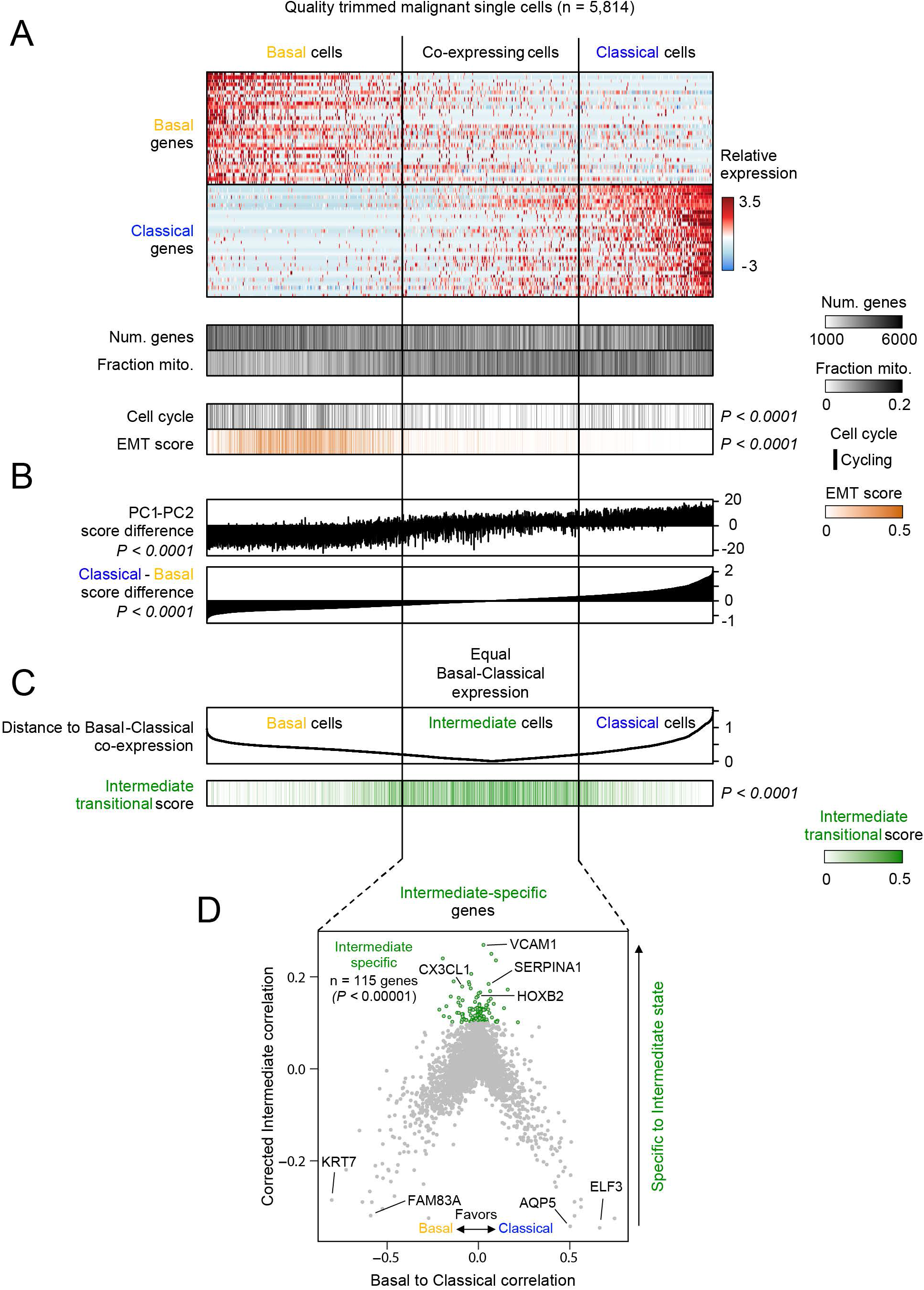
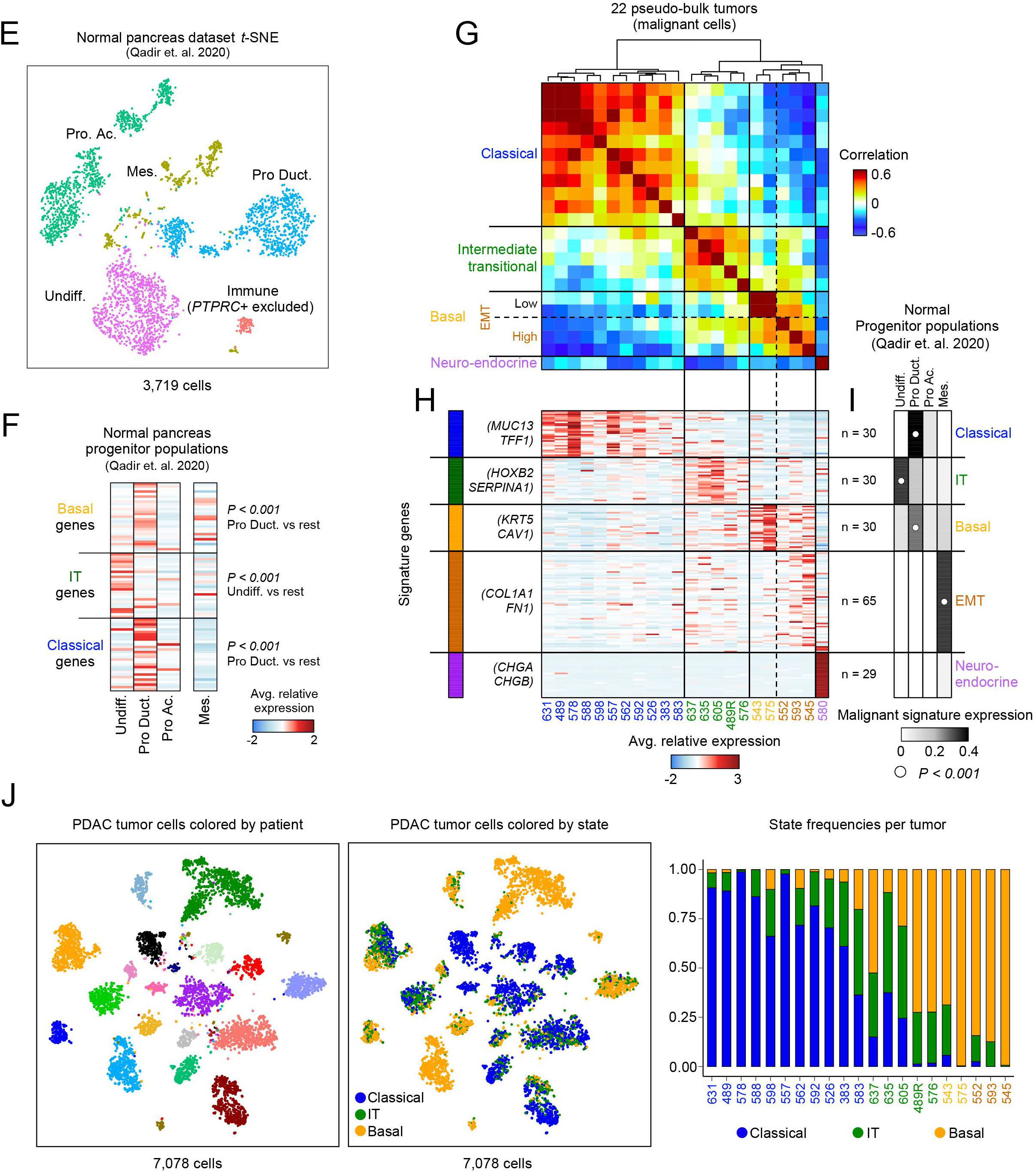
Cells with intermediate co-expressing phenotypes express a distinct gene program. *Related to Figure 2* **A**, Expression of basal and classical gene programs, with cells ordered by their basal-classical score difference. Quality metrics, EMT scores and the binarized cell cycle program are shown for each single cell below the heatmap. **B**, PC1 and 2 difference (top) and Classical – Basal score difference (bottom) are shown. Cells with equal basal and classical expression are associated with intermediate PC scores and cells are ordered as in **S3A**. **C**, Euclidian distance for each cell to co-expression (y = x) of basal (x) and classical (y) expression scores. Bottom track indicates the score derived from the genes specific to the intermediate state shown in **S3D** and explained in **Methods**. **D**, Gene correlation to either basal or classical score (x axis) or the corrected intermediate correlation (Euclidean distance in **S3C**, **Methods**). Green highlighted genes have corrected intermediate correlation >0.1 (*P<0.00001* above shuffled). *P*-value for binarized cycling group differences in **S3A** was calculated using Fisher’s Exact test. *P*-values for EMT score in **S3A** and group differences in **S3B** and **S3C** were calculated by Kruskal-Wallis test with multiple hypothesis correction. **E**, *t*-SNE visualization after dimensionality reduction and re-clustering for the normal progenitor populations identified in Qadir et al., 2020. Cell types are collapsed to those favoring Acinar (Pro Ac.), Ductal (Pro Duct.), or Undifferentiated (Undiff.) subsets. Mesenchymal cells (Mes.) are included as a non-epithelial reference and the small subset of immune cells was excluded from the comparisons. **F**, Averaged expression of all three malignant programs in normal pancreatic progenitor niche subsets and mesenchymal cells defined in Qadir et al., 2020. *P*-values for each set of genes are computed by Kruskal-Wallis test with multiple hypothesis correction. **G**, Pairwise correlation for biopsies with malignant cells (n = 22). Data are correlation coefficients for the average expression of all signature genes in the malignant cells from a given biopsy. EMT genes are from Groger et al., 2012. Clade identities are at left with the one PanNET tumor (PANFR0580) included for comparison and PANFR0604 not included due to lack of malignant cells captured. **H**, Average expression for the 184 genes used for clustering in **S3G**. Clade identity colors match text color in **S3G** and individual samples (columns) are ordered as in **S3G** and sample ID numbers are provided below. **I**, Scores for the expression of genes in **S3H** (grey scale heat) across the 4 main cell types found in the pancreatic progenitor niche (Qadir et al., 2020). White dot indicates the normal subset with the highest average expression for each malignant program (Kruskal-Wallis test), none of the normal subsets significantly express the Neuroendocrine gene signature. **J**, *t*-SNE visualization for malignant single cells in the biopsy cohort demonstrates intratumoral transcriptional heterogeneity at the single-cell level. Cells are colored by patient (left) or by transcriptional subtype (right).

**Supplemental Figure S4.**
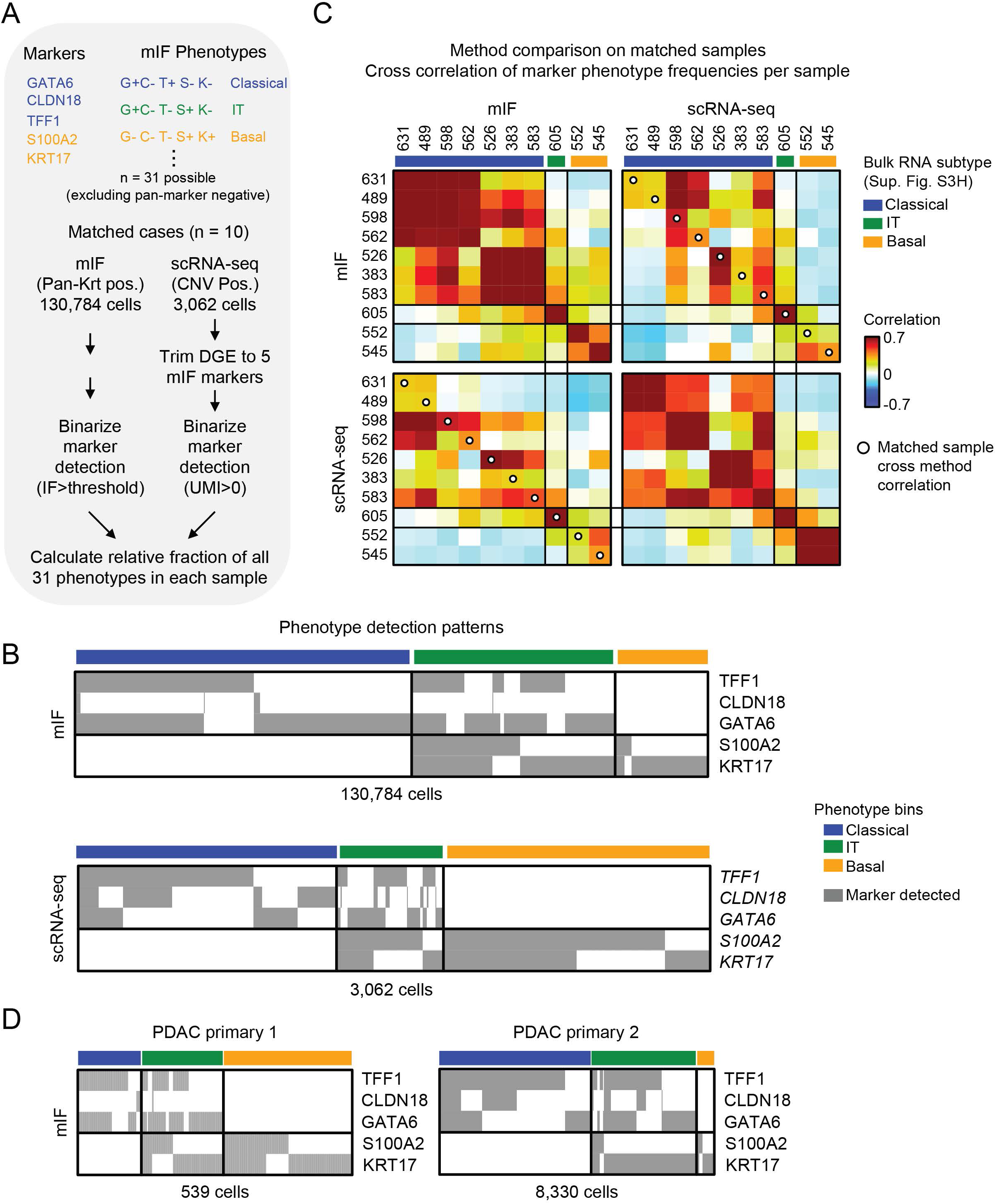
Multiplex immunofluorescence is concordant with scRNA-seq and demonstrates intratumoral heterogeneity with the presence of IT cells. *Related to Figure 2* **A**, Schematic for comparison of the matched datasets by combinatorial marker phenotypes. **B**, Marker detection in each single cell from the 10 samples in the mIF (top, 130,784 cells) and matched scRNA-seq datasets (bottom, 3,062 cells). Cells are sorted by their combinatorial phenotype outlined in **S4A**. **C**, Comparison within and between modalities on matched samples. Samples are sorted by the dendrogram in **Supplemental Figure S3G** and labeled with their pseudo-bulk RNA subtype identity. Correlation is performed over the fractional representation of each mIF phenotype (**S4A**) in each biopsy. Despite measuring different molecules (protein vs mRNA), the two approaches were highly concordant within RNA subtypes and on a case-by-case basis (white dots). **D**, mIF marker detection in each single cell from two primary PDAC samples shown in **Figure 2H**. Cells are sorted by their combinatorial phenotype outlined in **S4A**.

**Supplemental Figure S5.**
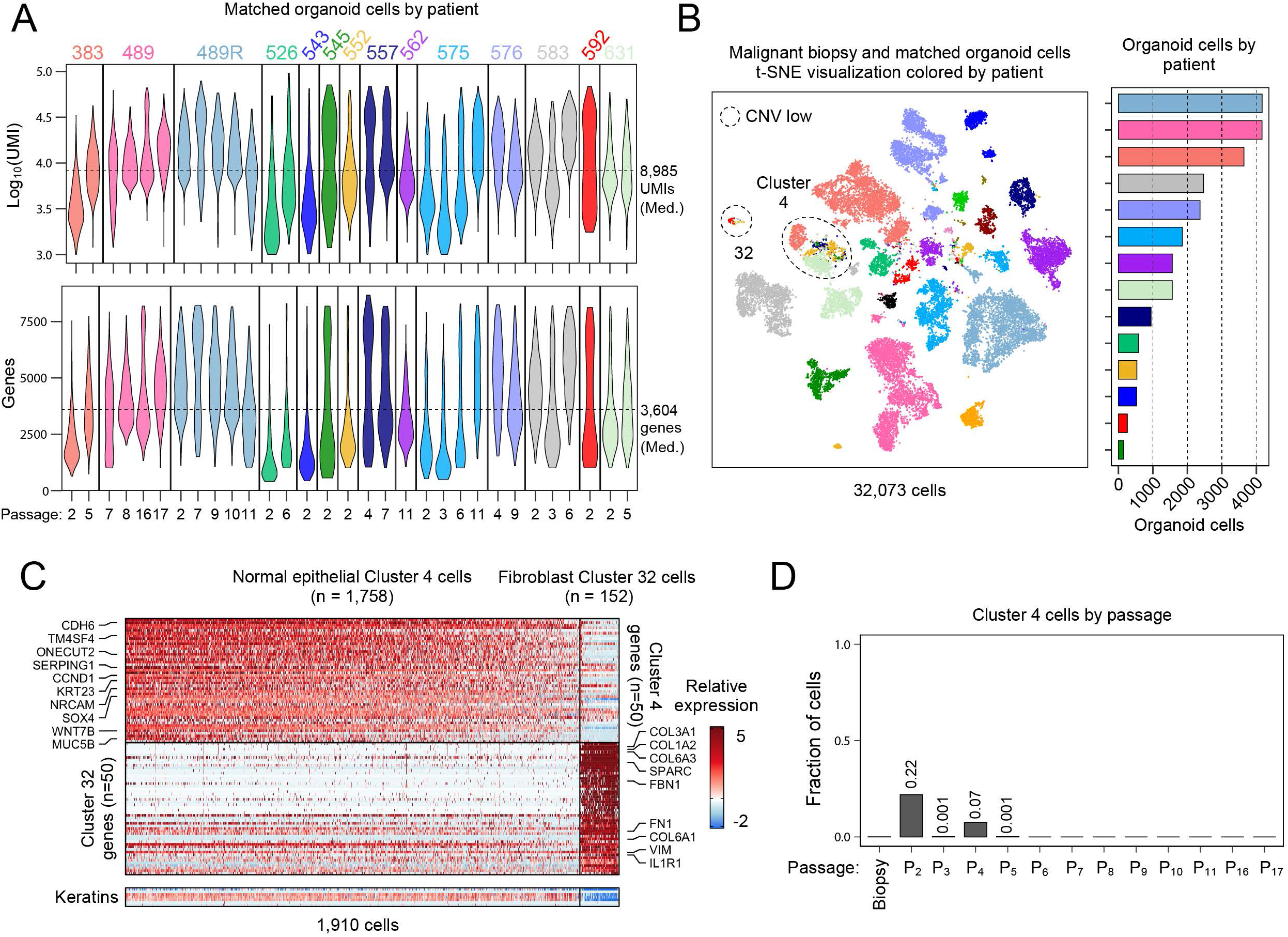
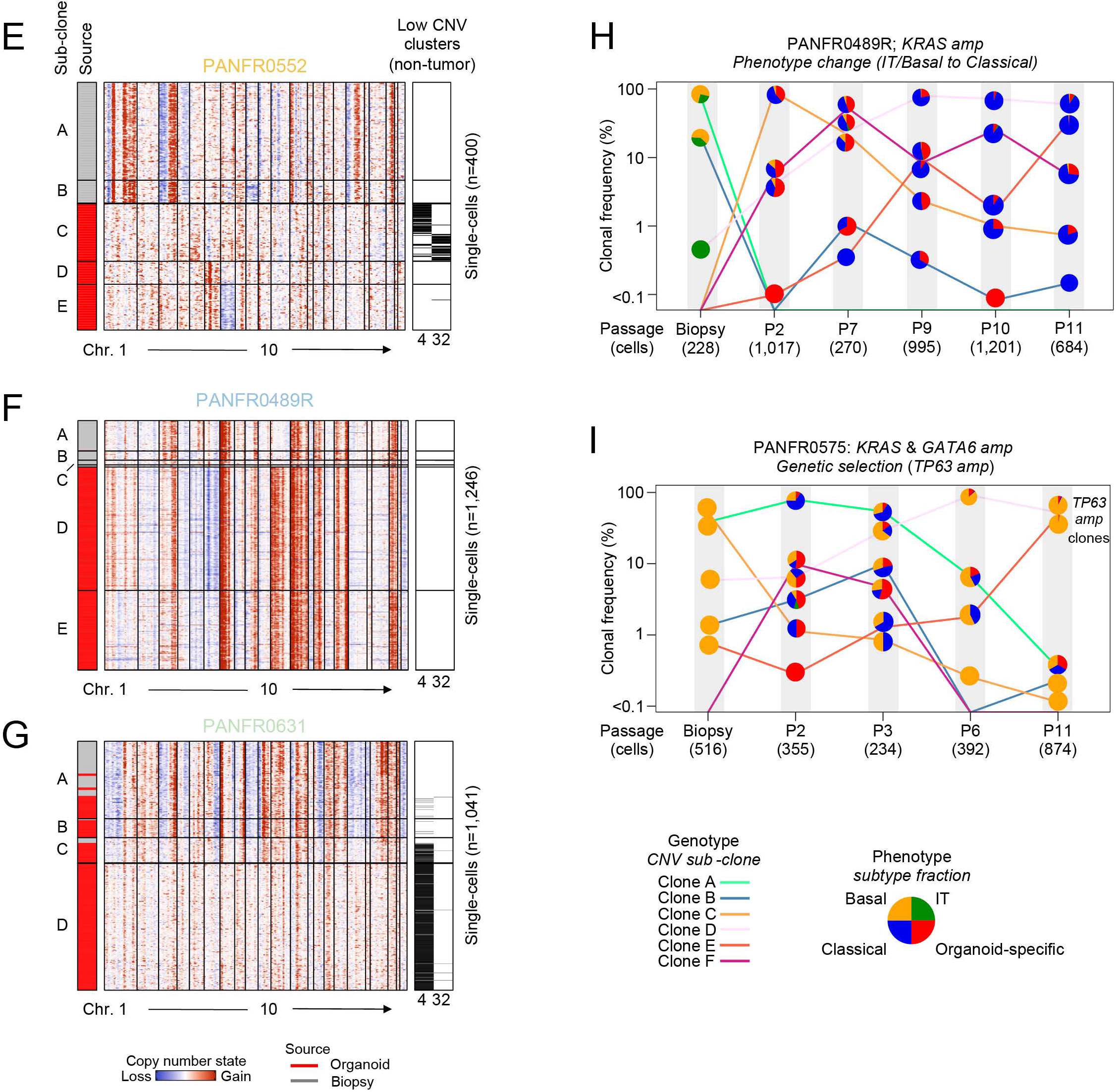
Quality metrics, cell type identification, and serial sampling across the patient-matched organoid cohort. *Related to Figure 4* **A**, Distribution of unique molecules and genes captured in quality cells per organoid sample, median values are indicated for each metric (dotted line) and violin plots are colored by patient ID (top, Log10(UMIs); bottom, number of genes). **B**, *t*-SNE visualization of all biopsy and matched organoid cells from iterative passages, colored by patient ID. Dotted circles indicate the only two SNN clusters (4 and 32) with appreciably admixed clusters and low CNV scores, the rest were patient-specific. Bar chart shows number of organoid cells recovered per matched sample (right). **C**, Relative expression for genes defining cluster 4 (top) and cluster 32 (bottom; 1 versus rest DE with the cells in **S5B**). Cluster 4 had an ambiguous epithelial identity while cluster 32 cells were defined by canonical fibroblast genes and low to absent detection of CNVs. **D**, Fraction of cluster 4 cells at each passage. These cells did not survive iterative passaging suggesting that they were either untransformed or unfit in organoid culture. **E-G**, Heatmaps show inferred CNV copy number status for every cell in each of three biopsy/early passage organoid pairs. Cells are ordered by hierarchical clustering of their CNV profiles and letters on the far left indicate subclones that have significant statistical evidence for tree-splitting (**Methods**). Each cell’s origin (biopsy tissue, grey; early passage organoid, red) is also noted (“Source” column). Right metadata bars indicate if that cell came from an admixed SNN cluster (4 or 32 in **S5B**). **H**, **I**, Matched phenotype and genotype evolution at each passage in PANFR0489R (**S5H**) and PANFR0575 (**S5I**). Frequencies of individual CNV clones at each time point (**Methods**, y axis) are tied by colored lines. Fill represents the transcriptional phenotype fraction for each CNV clone. In sample PANFR0575 (**S5I**), clones D and E had inferred *TP63* amplifications which expanded over time.

**Supplemental Figure S6.**
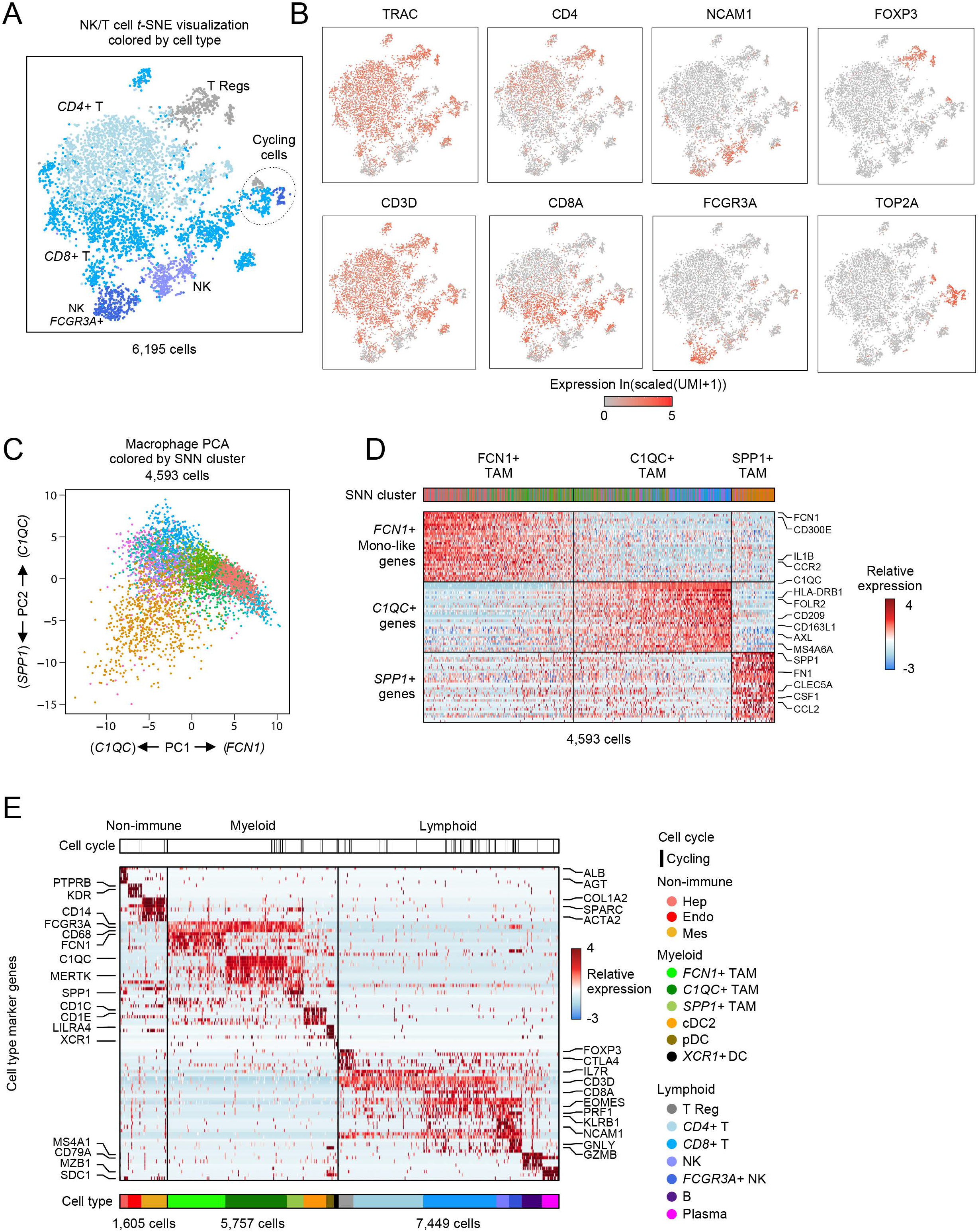
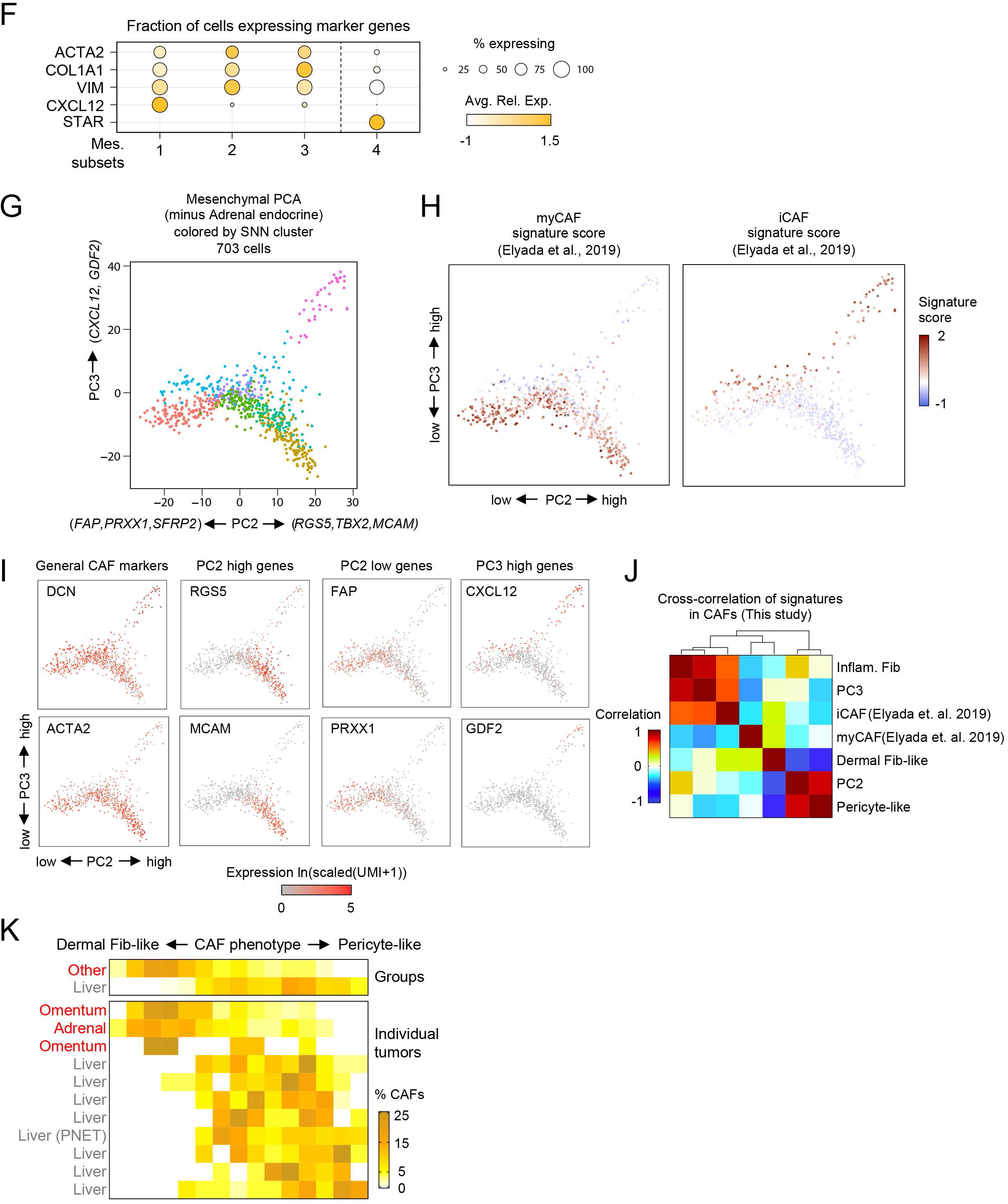
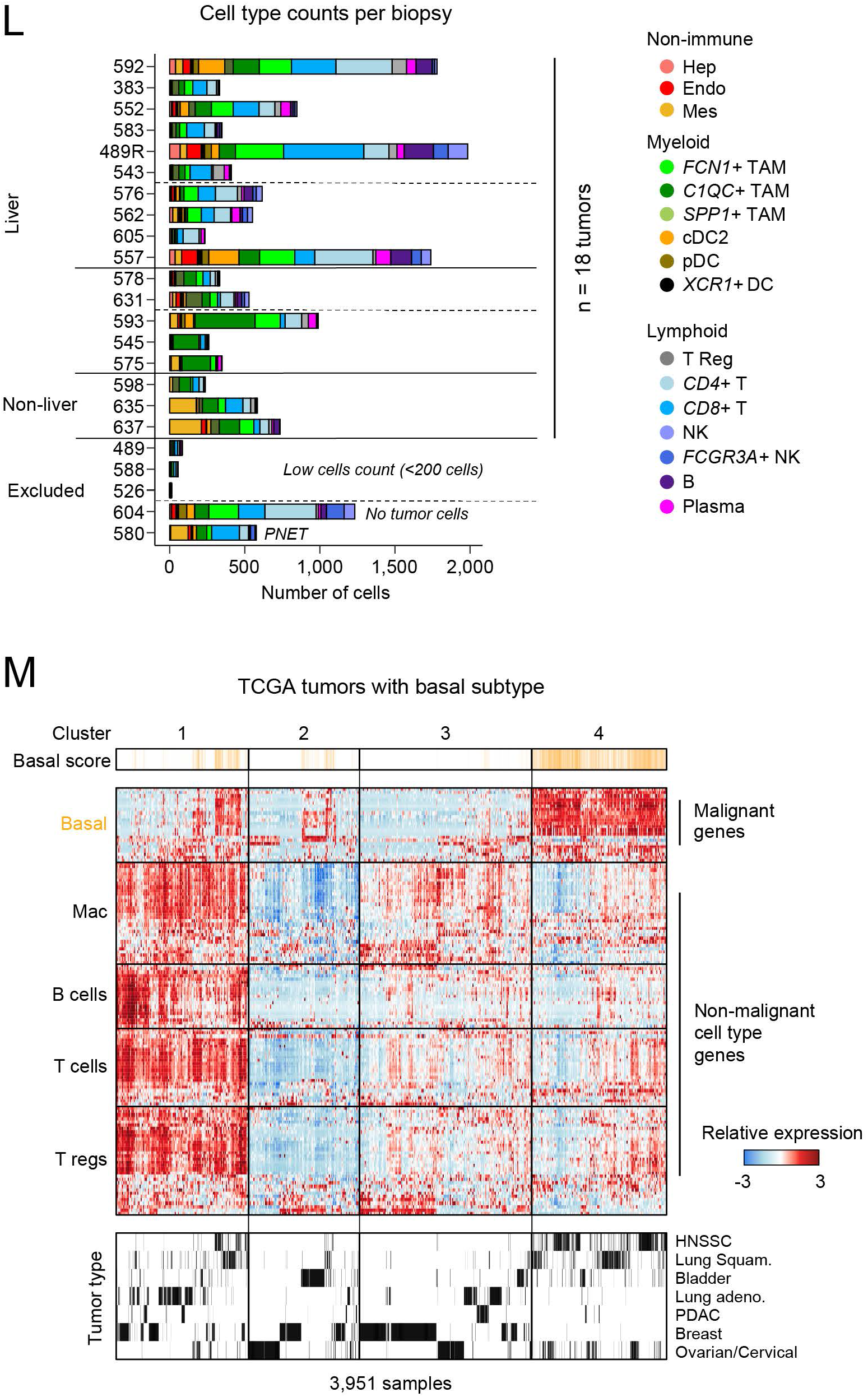
Identification of T/NK, macrophage, and fibroblast heterogeneity in the metastatic microenvironment. *Related to Figures 5 & 6* **A**, *t-*SNE visualization of sub-clustering (SNN) performed on T/NK cells in the metastatic cohort. Cells are colored by their type identity based on shared SNN cluster membership (Methods). **B**, Select cell type marker expression overlaid on the *t-*SNE visualization from **S6A**. **C**, PCA identifies 3 major subsets of TAMs in the metastatic niche. PC1 largely separates *FCN1*+ monocyte-like TAMs from more committed macrophage phenotypes. PC2 separates *SPP1*+ from *C1QC*+ macrophage phenotypes. **D**, Heatmap visualization of the gene expression programs specific to each TAM subset identified by the PCA in **S6C** (**Methods**). Top metadata indicate SNN cluster as in **S6C**. **E**, Heatmap shows the relative expression for select cell type markers. Top bar indicates the binarized cell cycle program (black, cycling) and the bottom color bar corresponds to the cell type colors noted in **Figure 6A**. **F**, Dot plots for average expression of the indicated CAF and adrenal endocrine marker genes in each of the cell subsets (1-4) identified in **Figure 5C**. Size of the dot indicates fraction of cells expressing a given gene. **G**, PCA over fibroblasts in the cohort (excluding Adrenal endocrine cells; subset 4, **Figure 5C**). Scatter plot of PC2 vs PC3 defines 3 states for CAFs in our cohort (**Methods**). **H**, Same visualization in **S6B**, but cells are colored by previously identified myCaf or iCaf signature scores. myCaf is evenly distributed across PC2 and iCaf associates with higher PC3 scores. **I**, Expression for select markers overlaid on the PCA from **S6B**. **J**, Cross-correlation of fibroblast signatures in single-cells. New dermal-vs. pericyte-like signatures provide non-overlapping information. PC3 inflammatory phenotypes are similar to the previously reported iCaf phenotype (Elyada et al., 2019) and our PC3-derived inflammatory fibroblast signature. **K**, Distribution across the CAF continuum comparing site differences as groups (top) or individual tumors (bottom). Heat indicates the fraction of CAFs in that score bin. **L**, Bar plot shows the number of non-malignant cells in each biopsy, color fill indicates the number of each cell type captured in that sample. Five biopsies were excluded from the analysis in **Figure 6A**-**C** because they either had low cell capture or were from a tumor with indeterminant malignant transcriptional subtype. Relevant samples are organized as in **Figure 6A**. **M**, Cross TCGA analysis for basal and immune cell type markers in epithelial tumors with known basal subtypes (Cancer Genome Atlas Research et al., 2013). Tumors with strong basal gene expression do not associate with strong immune infiltrates. Clusters were determined by dendrogram splitting and disease type for each sample is indicated below.

**Supplemental Figure S7.**
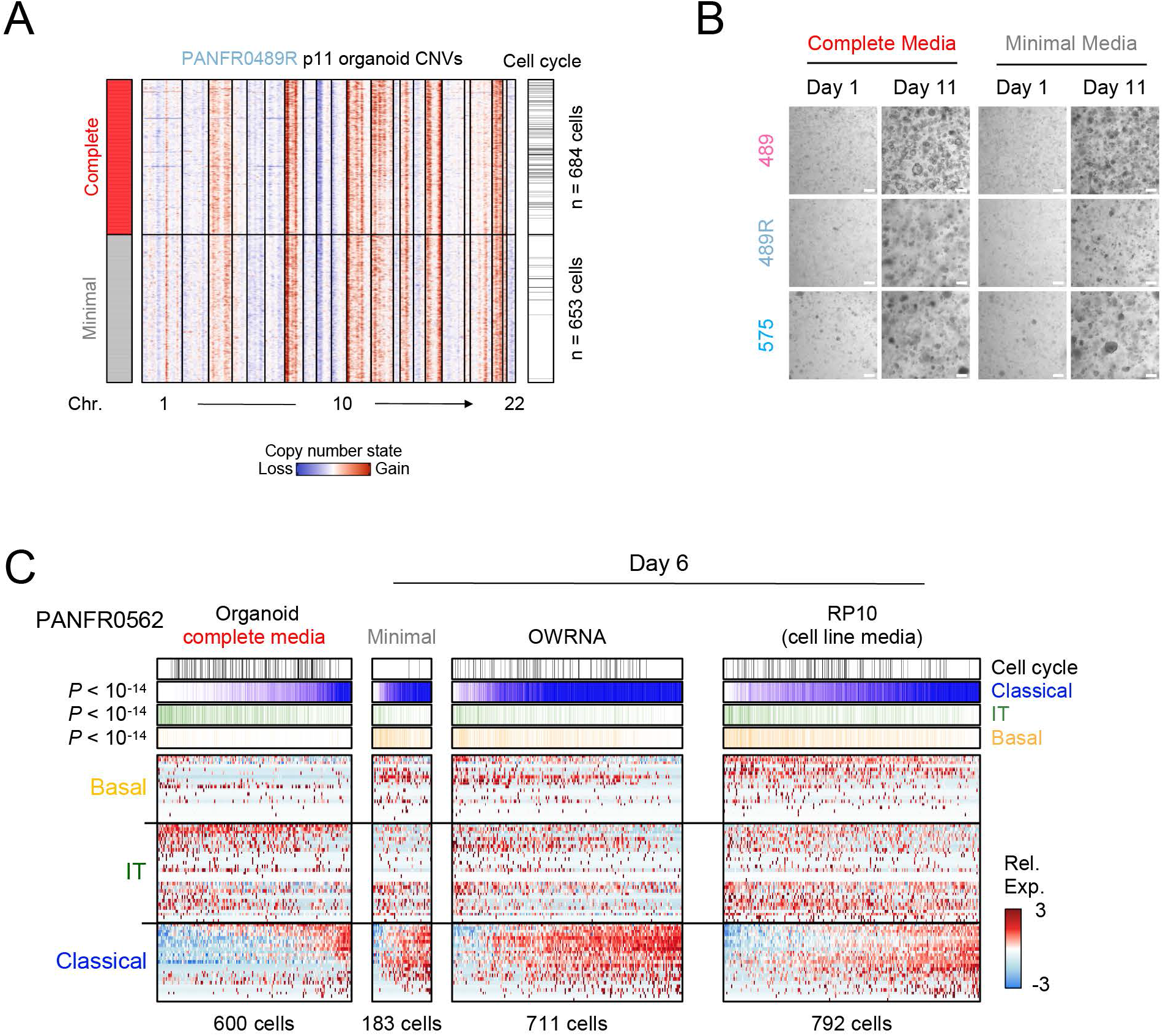
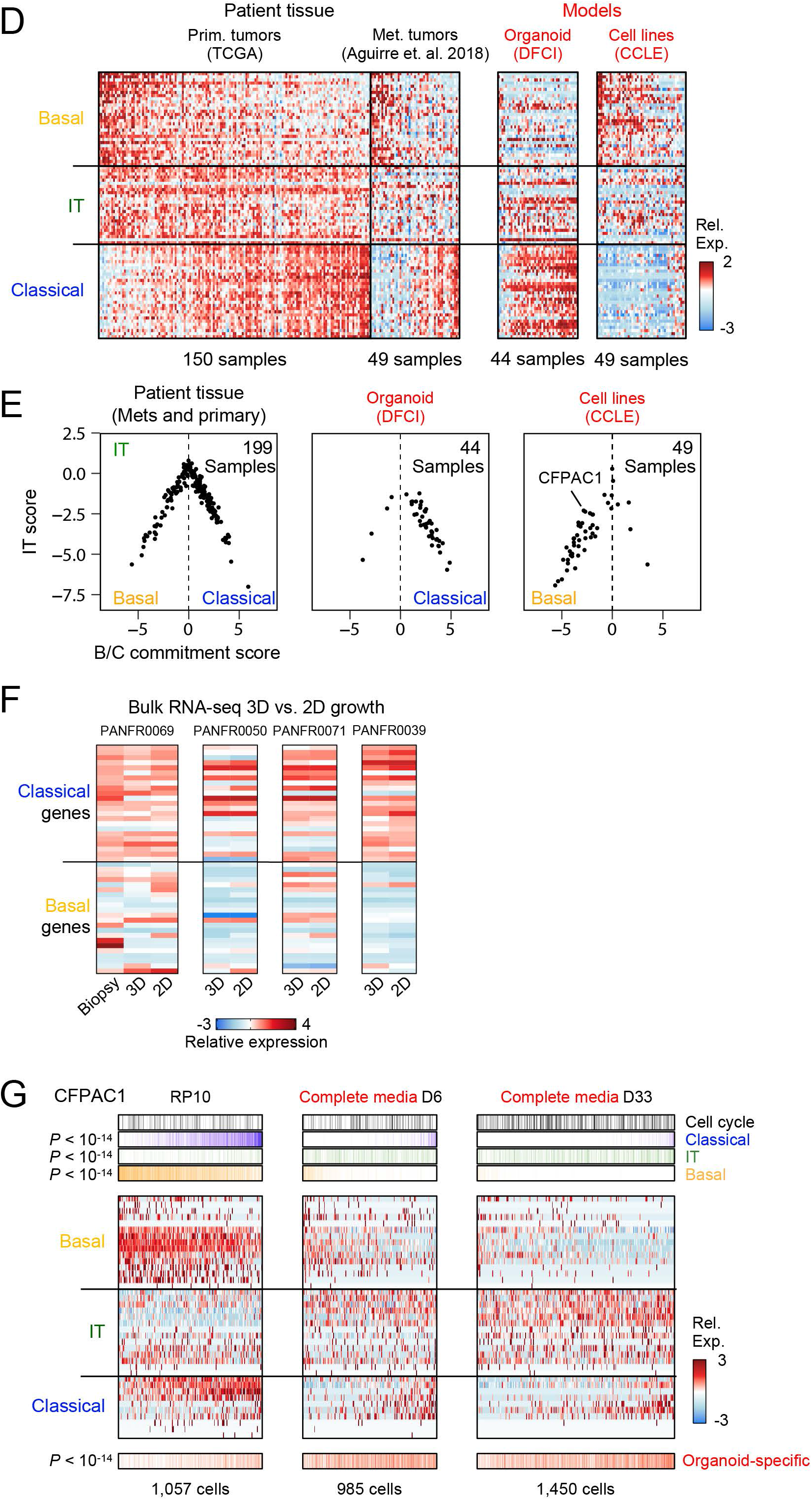
Alterations to organoid media, but not matrix dimensionality, shift transcriptional phenotype. *Related to Figure 7* **A**, Inferred CNVs for each cell from the PANFR489R samples cultured in either Minimal (grey) or Complete (red) organoid media conditions in **Figure 7B**. **B**, Brightfield images were obtained for organoids grown in standard organoid media (“Complete”) or in media without any growth factors (“Minimal”) at days 1 and 11 after seeding. **C**, Single organoid cells from model PANFR0562 (columns) cultured for 6 days in Complete medium, Minimal medium, OWRNA medium, or in RP10 (“cell line” medium, RPMI-1640 with 10% fetal bovine serum) and scored for basal, IT, and classical hierarchy phenotypes (rows). *P*-values for group differences were calculated by ANOVA followed by Tukey’s HSD. **D**, Relative expression for 90 genes representing PDAC state programs across bulk RNA-seq samples from primary resections (TCGA) and metastatic biopsies (Panc-Seq), as well as organoid and cell line (CCLE) models (Aguirre et al., 2018; Barretina et al., 2012; Cancer Genome Atlas Research Network, 2017; Ghandi et al., 2019). **E**, PDAC malignant state diagrams for average Basal-classical commitment score (x axis) and IT score (y axis) for bulk RNA-seq samples in **S7D**. **F**, Four established models were adapted to 2-dimensional culture in complete organoid media and measured via bulk RNA-seq. Rows indicate expression levels of basal-classical commitment score genes. **G**, Single cells from the established PDAC cell line CFPAC1 (columns) sampled in RP10 (standard “cell line” medium, RPMI-1640 with 10% fetal bovine serum) or at 2 timepoints in Complete organoid medium and scored for basal, IT, and classical phenotypes (rows). Bottom row indicates single cell organoid-specific gene expression (as described in **Figure 4B**) across all three conditions. *P*-values for group differences were calculated by ANOVA followed by Tukey’s HSD.

